# Identification of Key Genes and Underlying Mechanisms in Cystic Fibrosis and its Associated Complications Based on Bioinformatics Analysis of Next Generation Sequencing Data Analysis

**DOI:** 10.1101/2022.11.21.517323

**Authors:** Basavaraj Vastrad, Chanabasayya Vastrad

## Abstract

Cystic fibrosis (CF) is one of the inherited autosomal recessive disorders with very complicated pathogenesis. Identifying the molecular signatures and specific biomarkers of CF might provide novel clues for CF and and CF associated complications prognosis and targeted therapy. Based on the nexgeneration sequencing (NGS) dataset GSE136371 downloaded from the Gene Expression Omnibus (GEO) database, the differentially expressed genes (DEGs) between CF samples and normal controls were identified by using DESeq2 bioconductor package of R software. Gene ontology (GO) and REACTOME pathway enrichment analyses were applied for the DEGs. Then protein-protein interaction (PPI) network of these DEGs was visualized by Cytoscape with IMEx interactome database. The most significant module from the PPI network was selected for GO and pathway enrichment analysis. Subsequently, a miRNA-hub gene regulatory network and TF-hub gene regulatory network were constructed to identify hub genes, miRNAs and TFs. Finally, receiver operating characteristic curve (ROC) analysis was established to validate these hub genes. Total of 917 DEGs were identified between CF and normal control samples in GSE136371 dataset, including 479 up regulated and 438 down regulated genes. The most enriched DEGs in GO and pathway enrichment analysis were mainly associated with response to stimulus, regulation of cellular process, Neutrophil degranulation and rRNA processing. PPI network, module analysis, miRNA-hub gene regulatory network and TF-hub gene regulatory network predicted ten hub genes (FN1, UBE2D1, SRPK1, MAPK14, CEBPB, HSP90AB1, HSPA8, XRCC6, NCL and PARP1). In conclusion, the DEGs, relative pathways and hub genes identified in the present study might aid in understanding of the molecular mechanisms underlying CF and CF associated complications progression and provide potential molecular targets and biomarkers for CF and CF associated complications.

## Introduction

Cystic fibrosis (CF) is a life-threatening inherited autosomal recessive disorder and affects at least 100 000 people worldwide [1]. In patients with CF, quality of life especially in children and adolescents is seriously affected by severe damage to the lungs, digestive system and other organs in the body [2]. Pancreatitis [3], gastrointestinal complications [4], chronic endobronchial airway infection [5], pulmonary complications [6], sinusitis [7], diabetes mellitus [8], nasal polyps [9], antibiotic resistance [10], depression and anxiety [11], liver disease [12], osteoporosis [13], bronchiectasis [14] and arthritis [15], remain the most common complications for patients with CF. CF is characterized by pancreatic insufficiency and chronic endobronchial airway infection [16]. At current, there is no effective drug for treatment of the CF [17], and, because prenatal diagnosis is the most effective way to avoid the birth of children with CF, it is essential to investigate the molecular pathogenesis of this disease.

The fast development of next generation sequencing technology and bioinformatics analysis based on high-throughput data, provide novel tactics to identify differentially expressed genes (DEGs) and discover therapeutic targets for the initiation and evolution of CF [18]. Altered expression of genes and signaling pathways plays an important role in the initiation and progression of CF. Recently, studies identified altered expression of genes include ADIPOQ (adiponectin, C1Q and collagen domain containing) and STATH (statherin) [19], WISP1 [20], TAS2R38 [21], STAT3, IL1B and IFNGR1 [22] and CFTR (cystic fibrosis transmembrane conductance regulator) [23] were involved in progression of CF. Signaling pathways include S-nitrosothiols signaling pathway [24], MAPK signaling pathway [25], protein kinase signaling pathway [26], TGF-B signaling pathways [27] and IL-1 signaling pathway [28] were associated with progression of CF. In sum, it still remains unclear, more investigations are needed to understand the molecular pathogenesis in CF.

Prediction results of next generation sequencing (NGS) and bioinformatics analysis have become increasingly useful in the diagnosis and therapy of various diseases [29]. In addition to being used to identify the functional connections among genes in an unbiased manner for the in-depth investigations of biological processes, it can also be used to predict genes and to explore the relationship between gene expression and CF [30].

In this investigation, we performed a biological information analysis using NGS data GSE136371 [31] was downloaded from Gene Expression Omnibus database (GEO, https://www.ncbi.nlm.nih.gov/geo/) [32] and identified the DEGs for the CF and normal control samples. Subsequently, the Gene Ontology (GO), REACTOME pathway enrichment analysis, protein-protein interaction (PPI) network, modules, miRNA-hub gene regulatory network and TF-hub gene regulatory network were analyzed to understand the molecular mechanisms underlying CF. The diagnostic value of the identified hub genes was assessed by receiver operating characteristic curve (ROC) analysis. In conclusion, our investigation aimed to explore the molecular biomarkers of CF based on bioinformatics analysis and provide candidate biomarkers for early diagnosis and therapeutic targets.

## Materials and Methods

### Next generation sequencing data source

The NGS data, GSE136371 [31], from the GEO database was downloaded. GSE136371 is based on the GPL20301 Illumina HiSeq 4000 (Homo sapiens) and contains a total of 49 samples, including 33 CF samples and 19 normal control samples.

### Identification of DEGs

The R package, “DESeq2” [33] was used to analyze the GSE136371 NGS data. The Benjamini–Hochberg method was used to adjust original p-values, and the false discovery rate (FDR) procedure was used to calculate fold changes (FC) [34]. The differential gene expression threshold was log2 FC > 0.635 for up regulated genes, log2 FC < −0.12 for down regulated genes and adj p-value < 0.05. The ggplot2 package was used to visualize the DEGs into a volcano map, while the gplot package was used to cluster the significant DEGs.

### GO and pathway enrichment analyses of DEGs

The Gene Ontology (GO, http://www.geneontology.org) [35] is a bioinformatics resource. It provides information about genes and gene product functions and exploits ontology to strengthen biological knowledge. The REACTOME (https://reactome.org/) [36] is a database for the qualitative interpretation of genomic sequences and other biological data, including systematic, genomic, and chemical information as well as an supplementary human specific class of health information. GO enrichment analyses was performed in terms of biological processes (BP), cellular components (CC), and molecular function (MF). The related biological functions and signal pathways were analyzed using GO/ REACTOME enrichment and analyzed again with the g:Profiler (http://biit.cs.ut.ee/gprofiler/) [37], with p < 0.05 considered to be statistically significant.

### Construction of the PPI network and module analysis

To investigate the protein interactions of DEGs, we submitted them to the IMEx interactome database (http://www.imexconsortium.org/) [38]. We then used Cytoscape software (http://www.cytoscape.org/) [39] (version 3.9.1) to integrate and visualize the PPI network. In addition, hub genes in the network were identified using the Network Analyzer application in Cytoscape software. The maximum group centrality of each gene in the network was calculated by the maximal node degree [40], betweenness [41], stress [42] and closeness [43] score. PEWCC1 [44] was applied to screen the modules of the PPI network, and the core modules were selected.

### miRNA-hub gene regulatory network construction

Interactions between miRNAs and hub genes were predicted using miRNet database (https://www.mirnet.ca/) [45], which integrated the prediction results of TarBase, miRTarBase, miRecords, miRanda (S mansoni only), miR2Disease, HMDD, PhenomiR, SM2miR, PharmacomiR, EpimiR, starBase, TransmiR, ADmiRE, and TAM 2. Cytoscape software [39] (version 3.9.1) was used to visualize the regulatory network.

### TF-hub gene regulatory network construction

Interactions between TFs and hub genes were predicted using NetworkAnalyst database (https://www.networkanalyst.ca/) [46], which integrated the prediction results of JASPER. Cytoscape software [39] (version 3.9.1) was used to visualize the regulatory network.

### Receiver operating characteristic curve (ROC) analysis

The drawing of the ROC curves and the calculation of the area under the curve (AUC) were conducted by the ‘pROC’ package in R [47]. Thus, we studied the feasibility of the hub genes for prediction using the AUC value. An area under the curve (AUC) > 0.8 indicated a good diagnostic value.

## Results

### Identification of DEGs

Based on the DEG selection criteria log2 FC > 0.635 for up regulated genes, log2 FC < −0.12 for down regulated genes, with adj p-value < 0.05), a total of 917 DEGs between CF samples and normal control samples were identified (Table 1), and comprising 479 up regulated genes and 438 significantly down regulated genes was selected for subsequent analysis. After screening the differential genes based on the volcano plot and cluster analysis were carried out, as shown in Fig. 1 and Fig. 2.

**Fig. 1.**
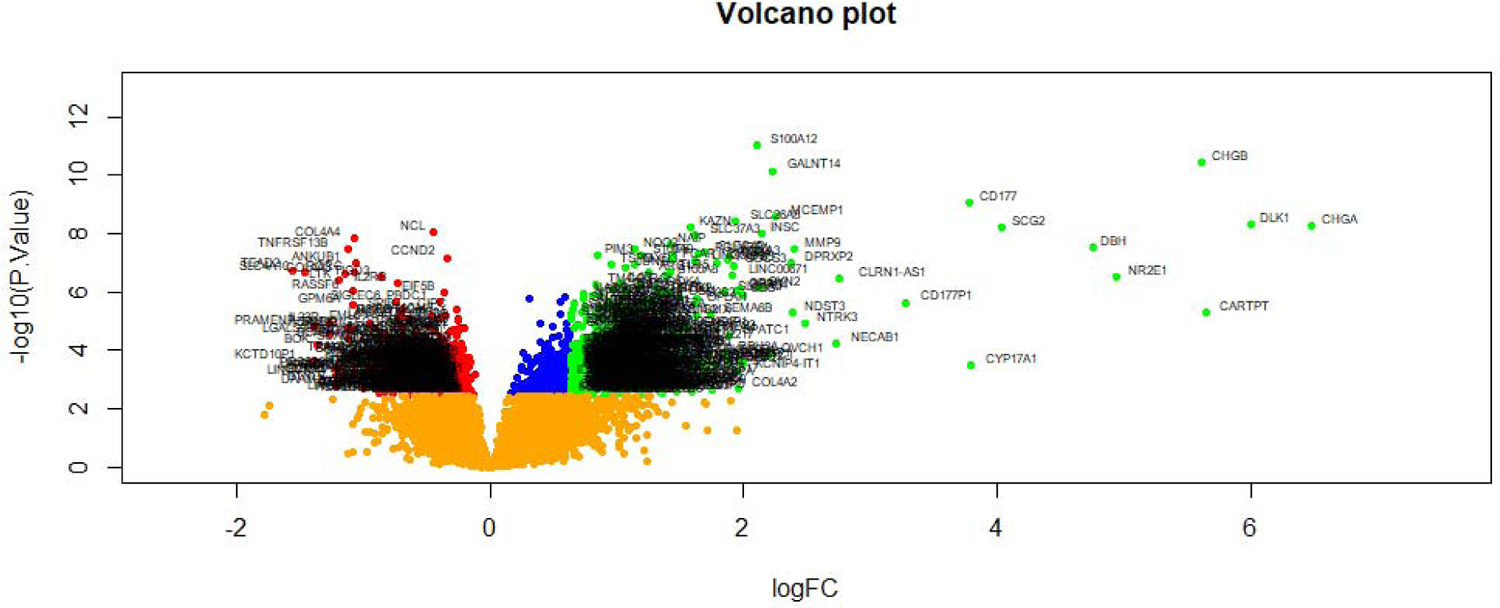
Volcano plot of differentially expressed genes. Genes with a significant change of more than two-fold were selected. Green dot represented up regulated significant genes and red dot represented down regulated significant genes.

**Fig. 2.**
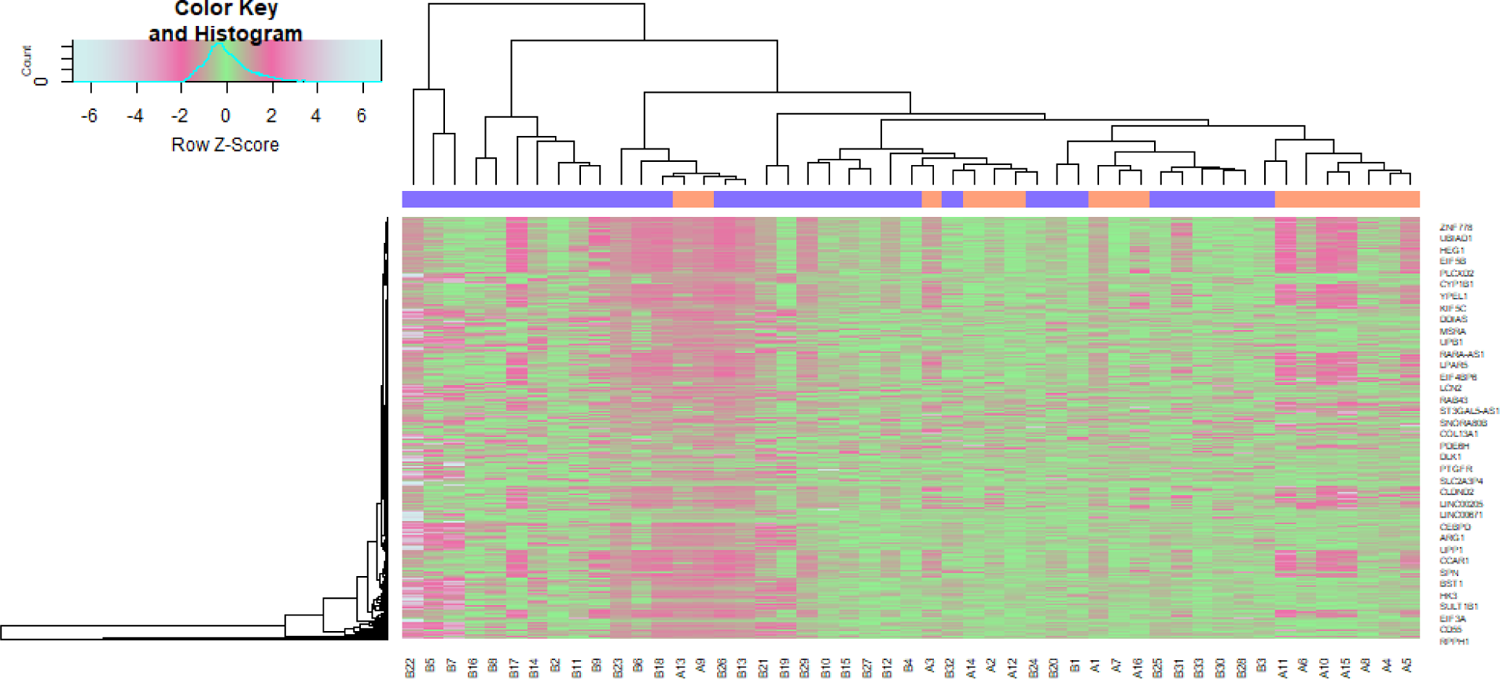
Heat map of differentially expressed genes. Legend on the top left indicate log fold change of genes. (A1 – A19 = normal control samples; B1 – B33 = CF samples)

**Table 1.**
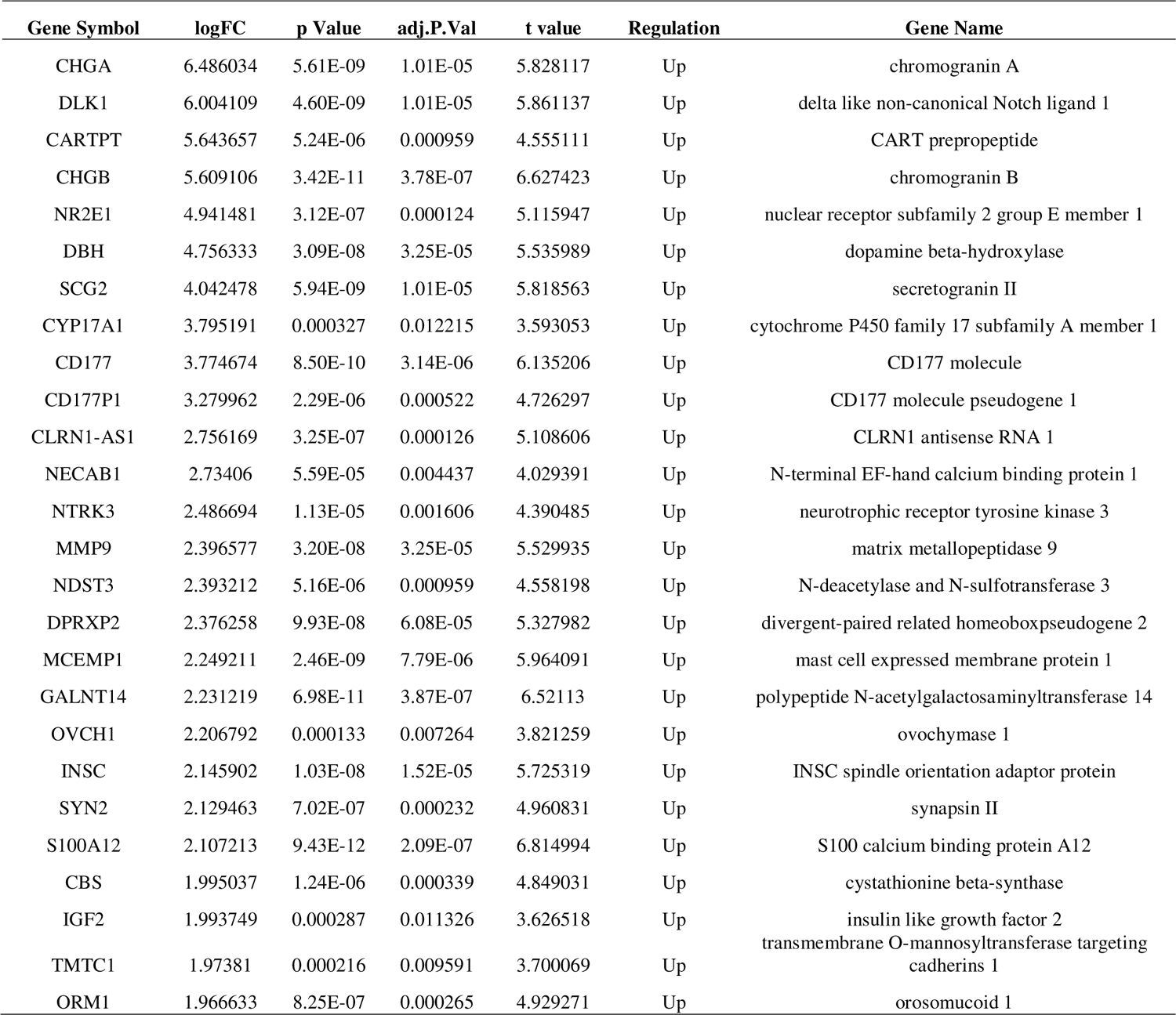
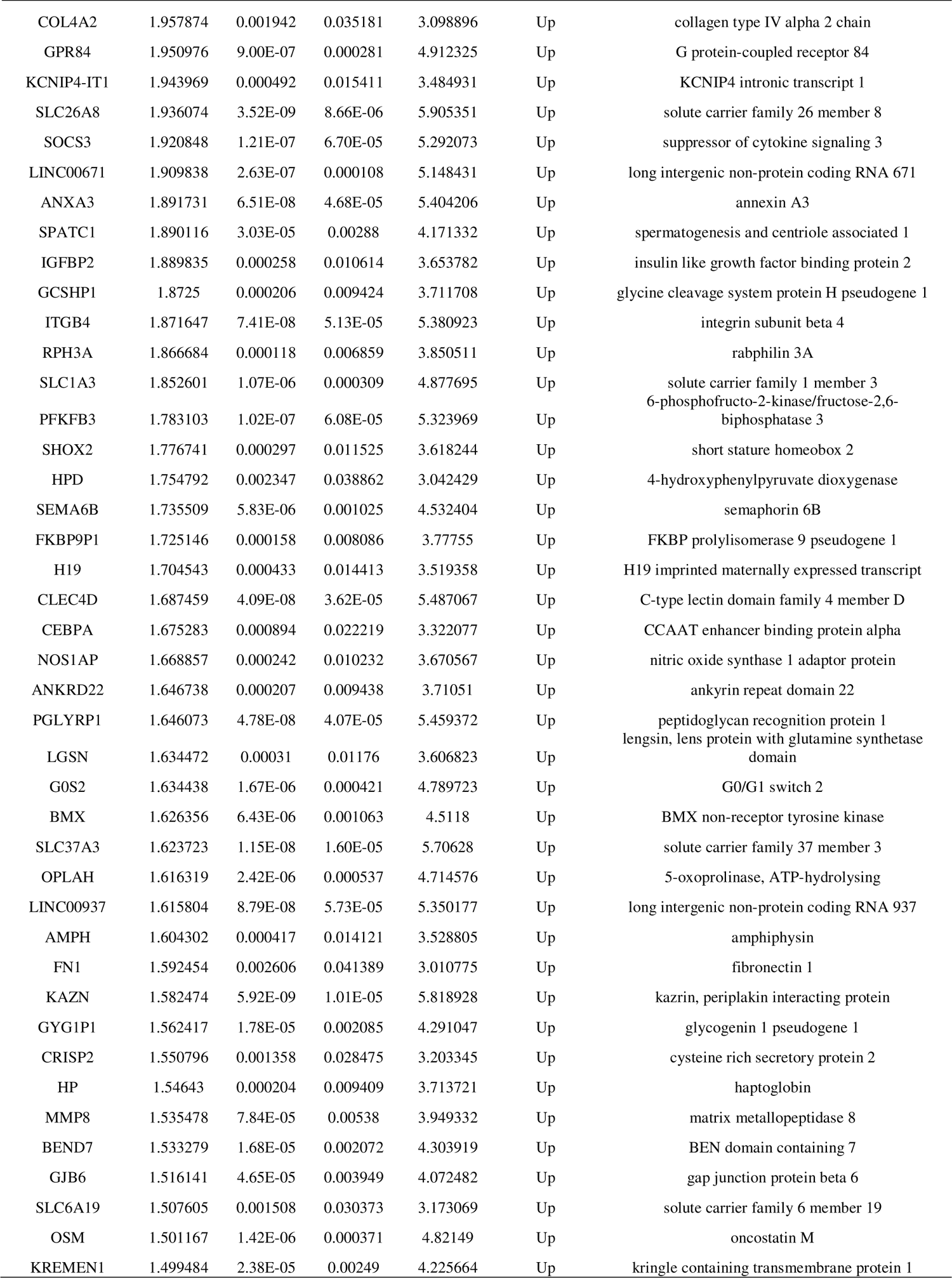
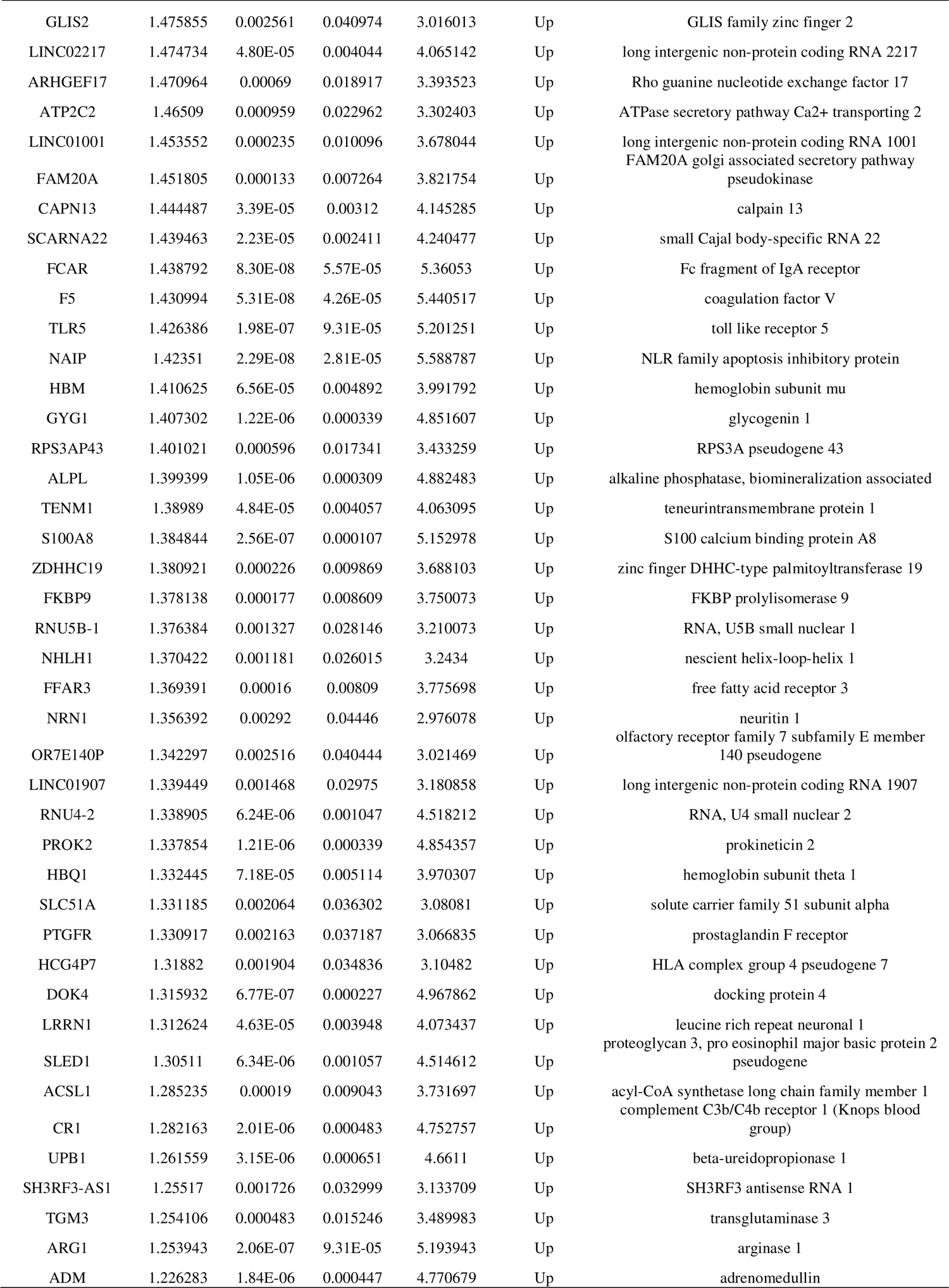
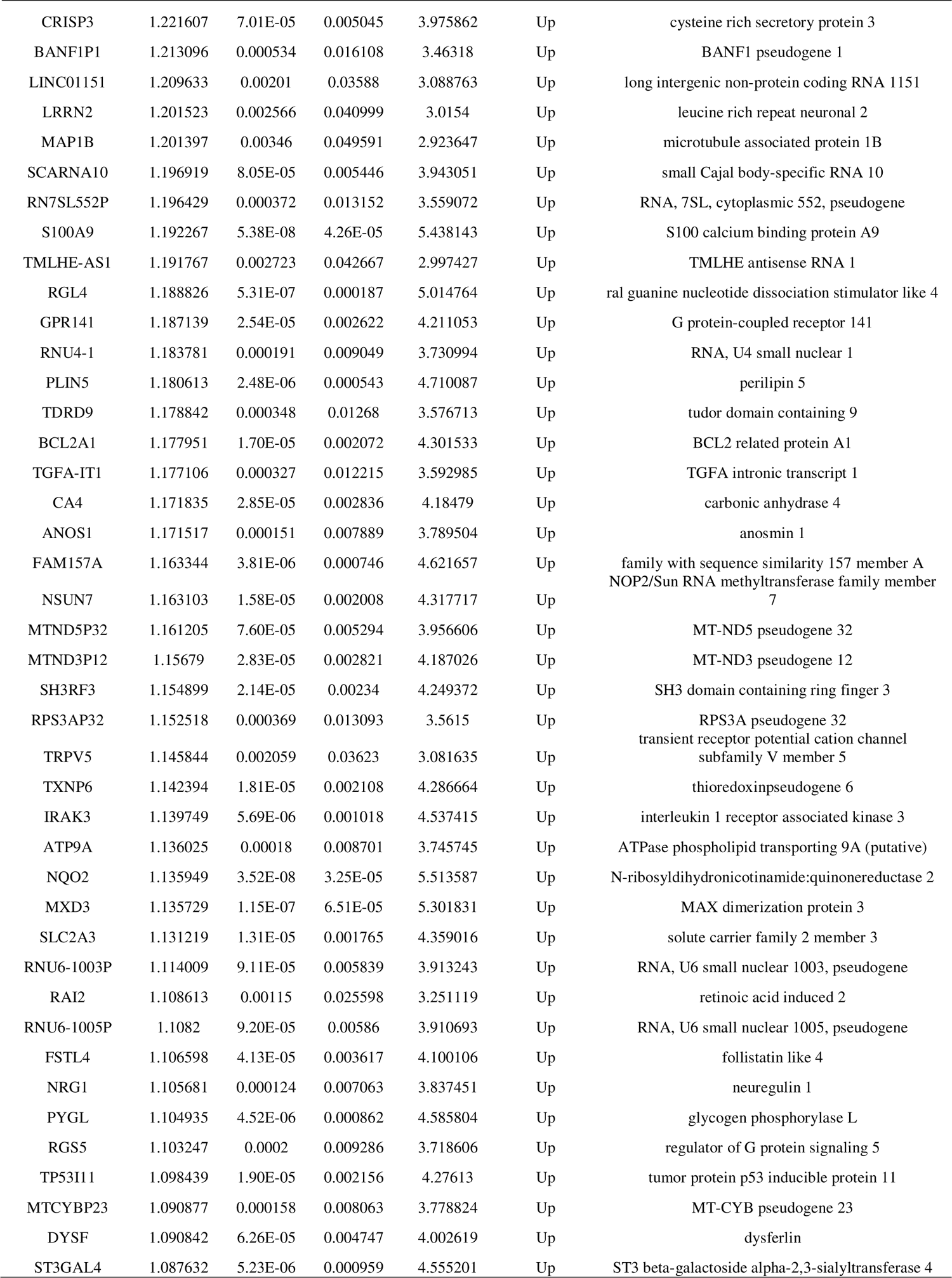
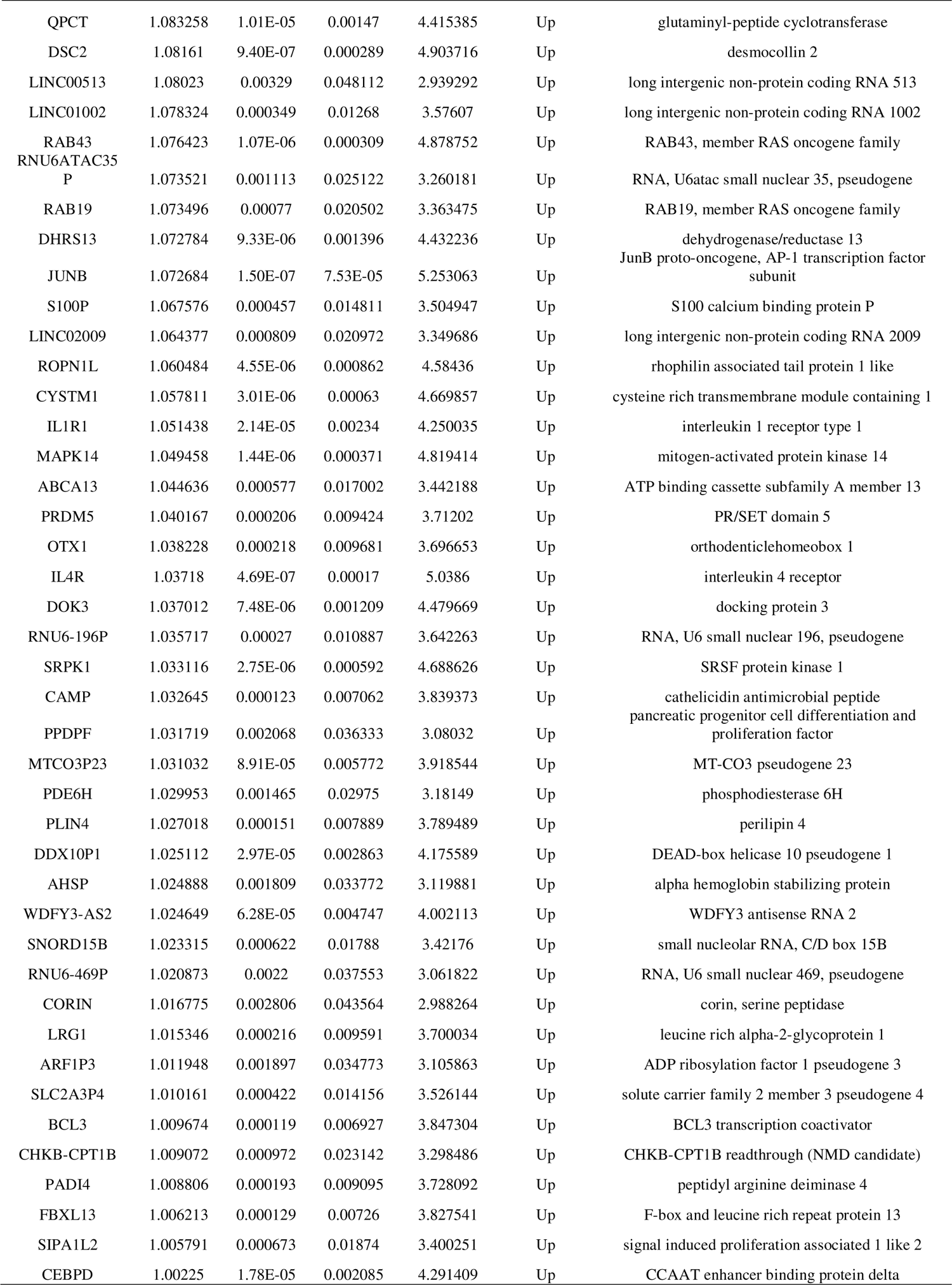
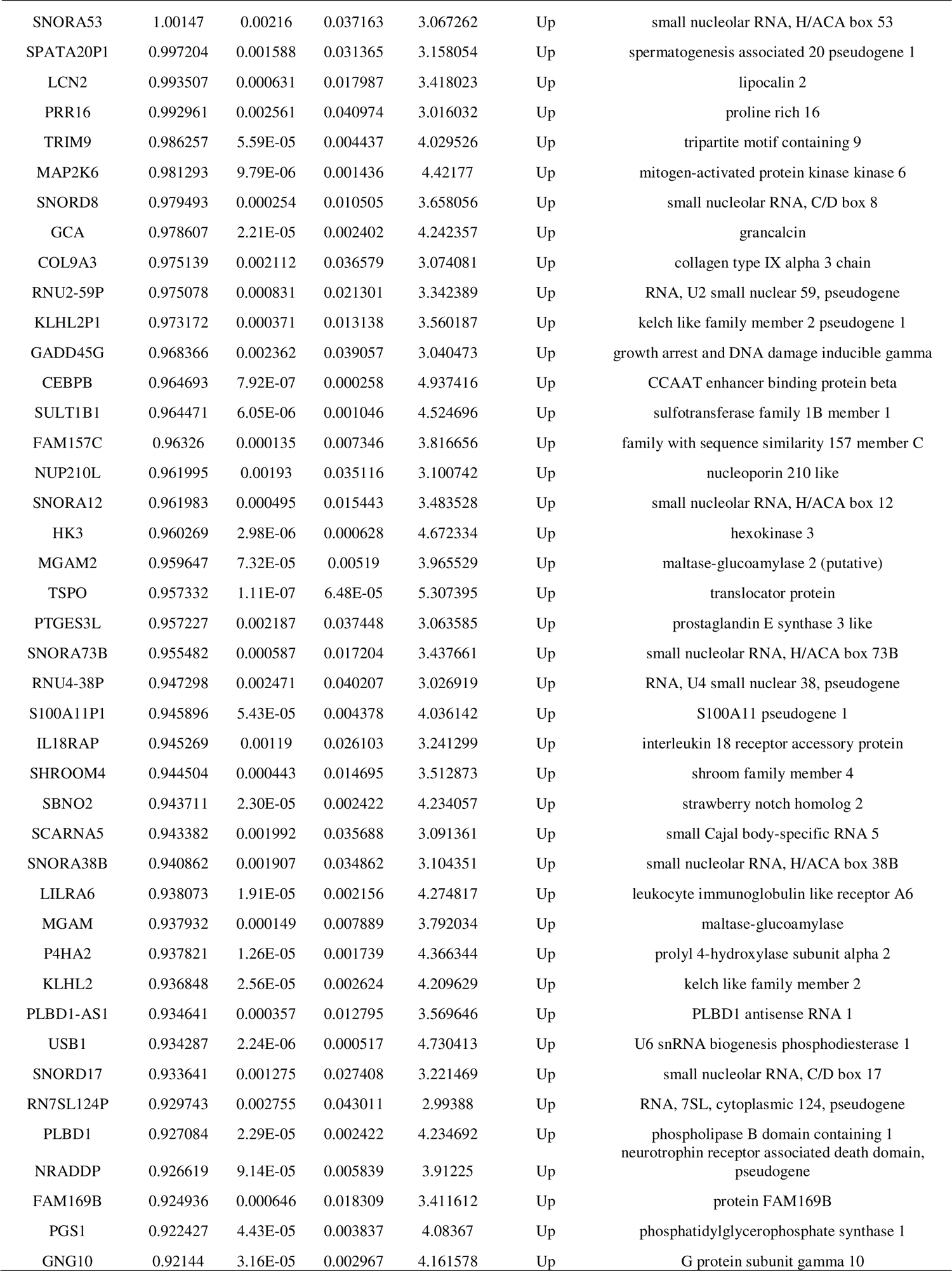
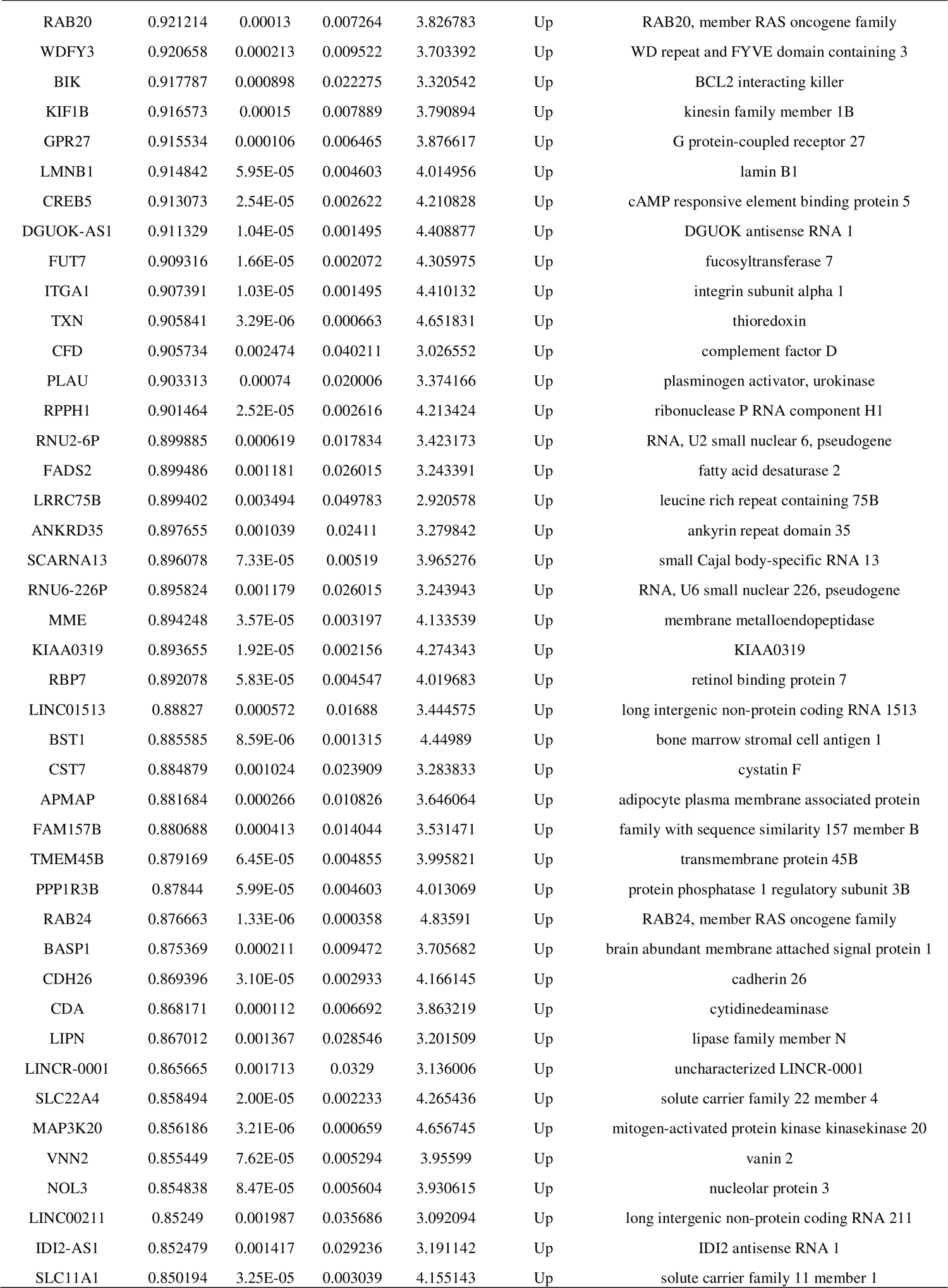
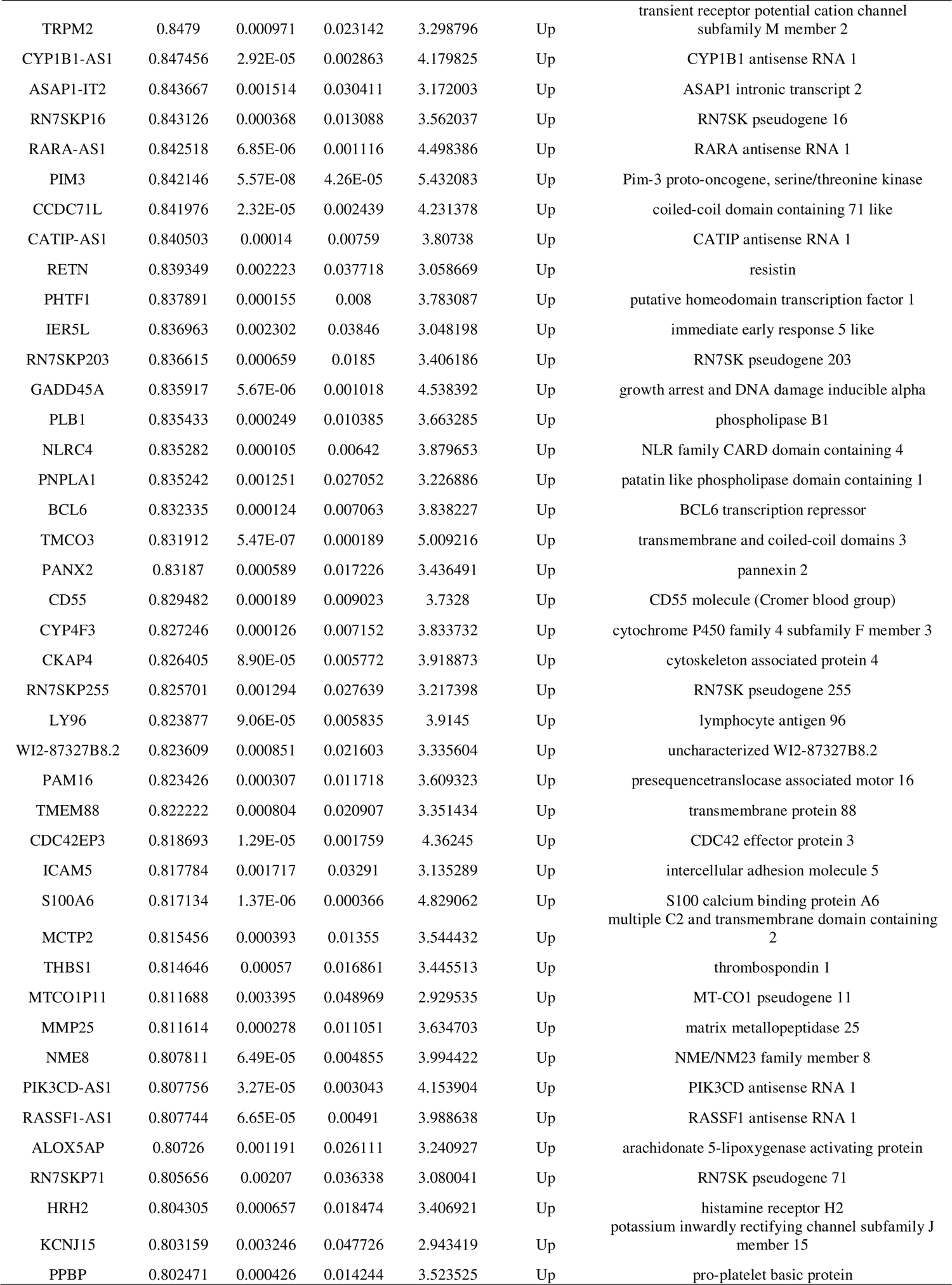
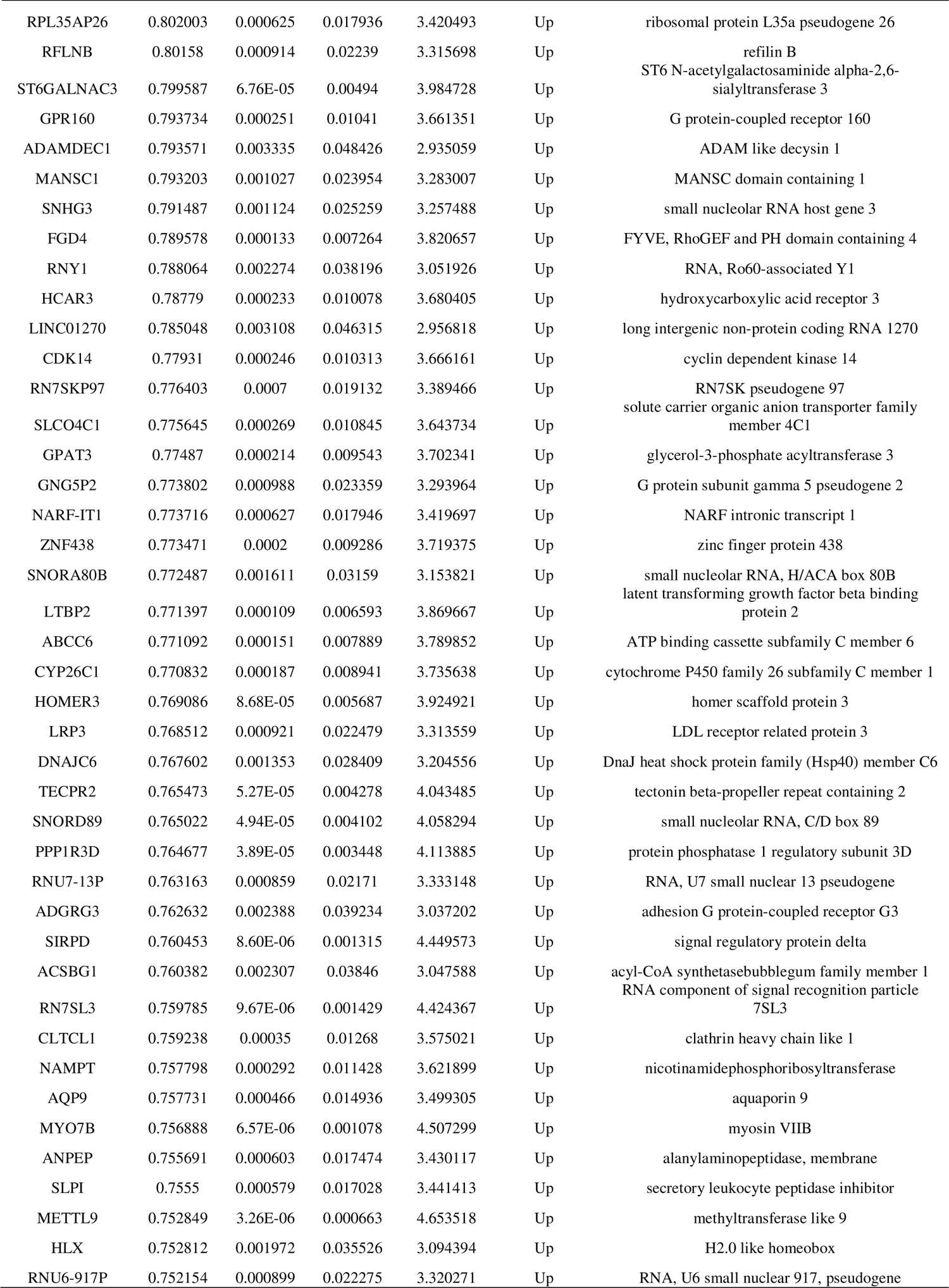
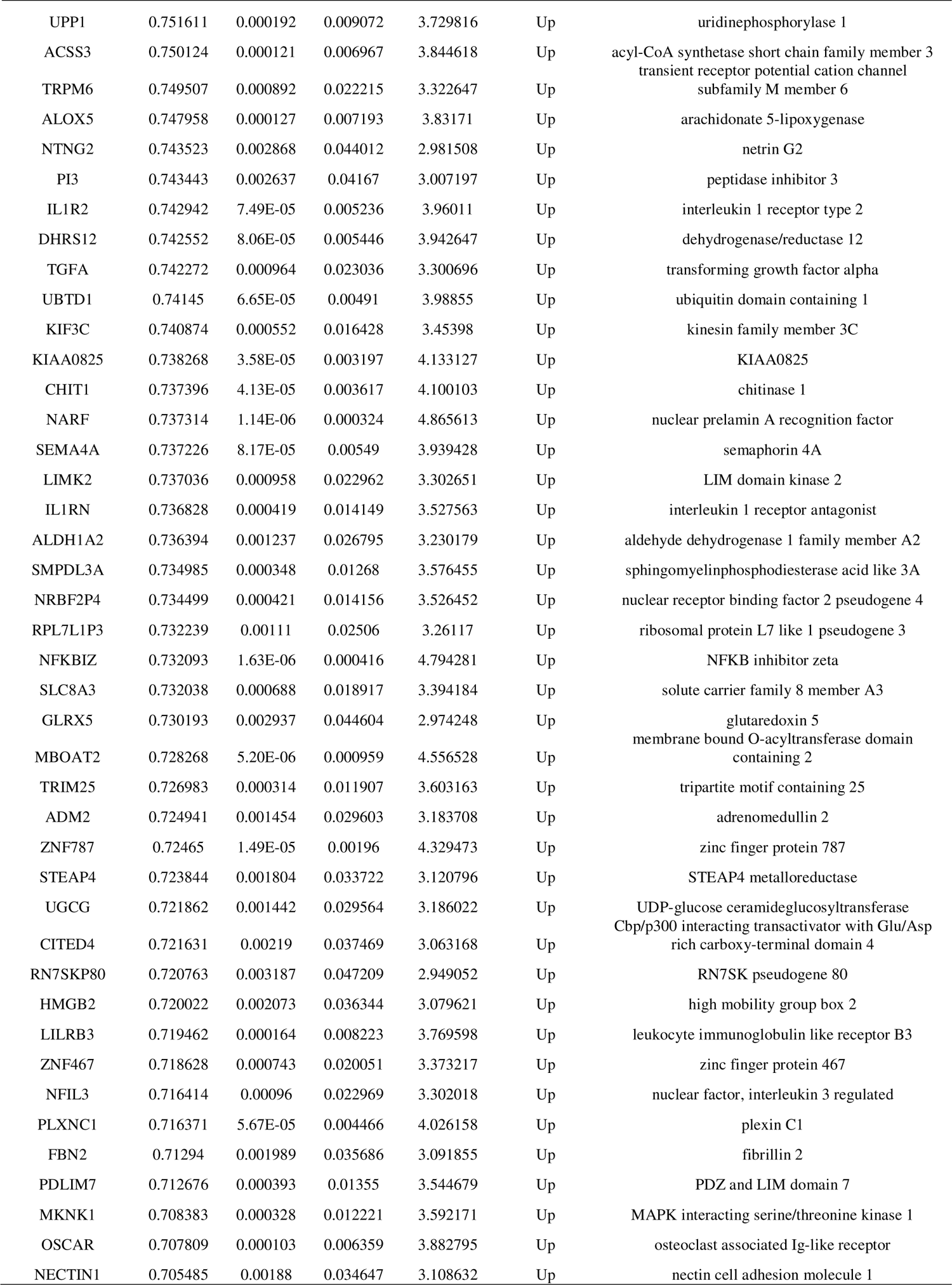
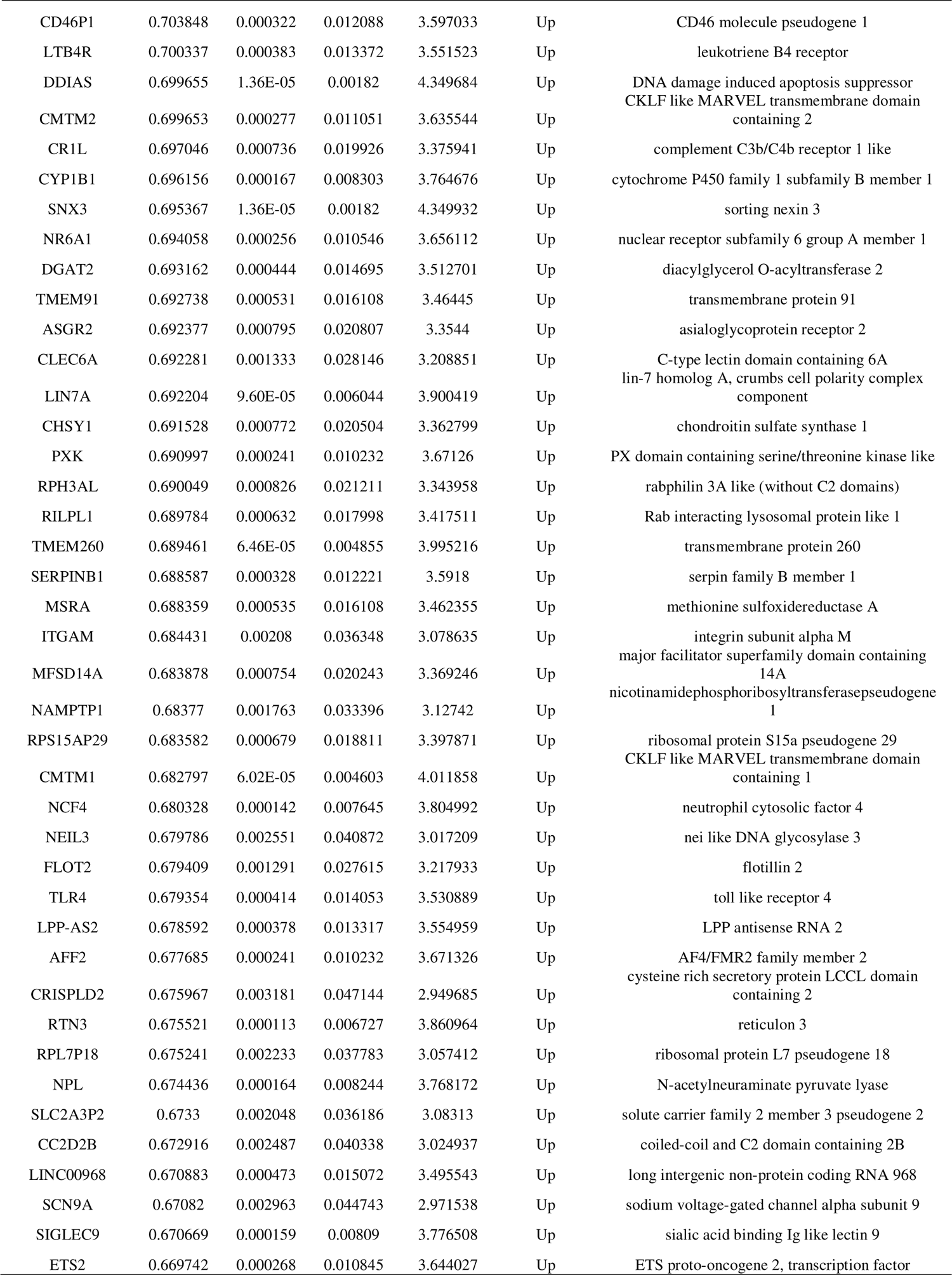
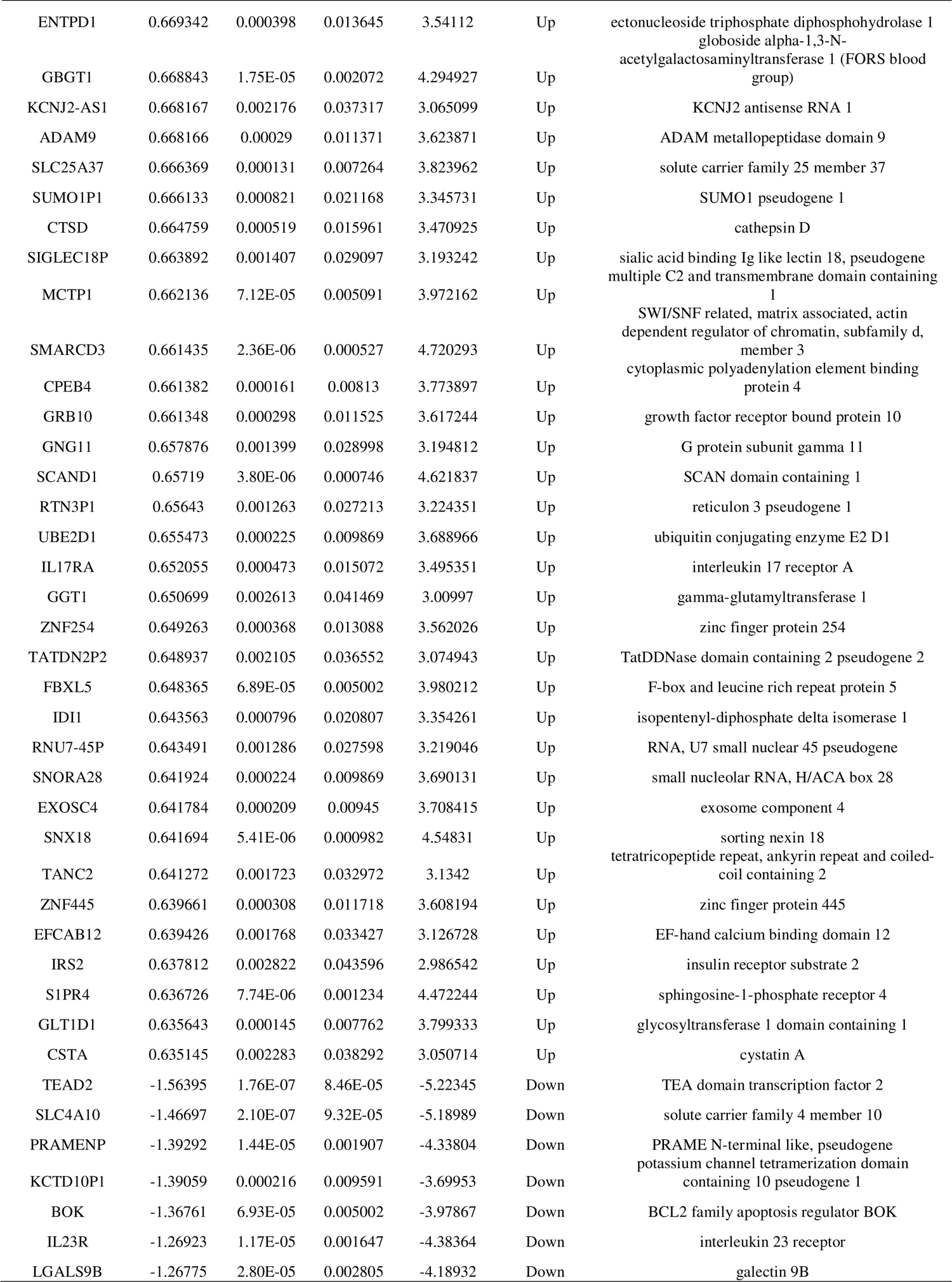
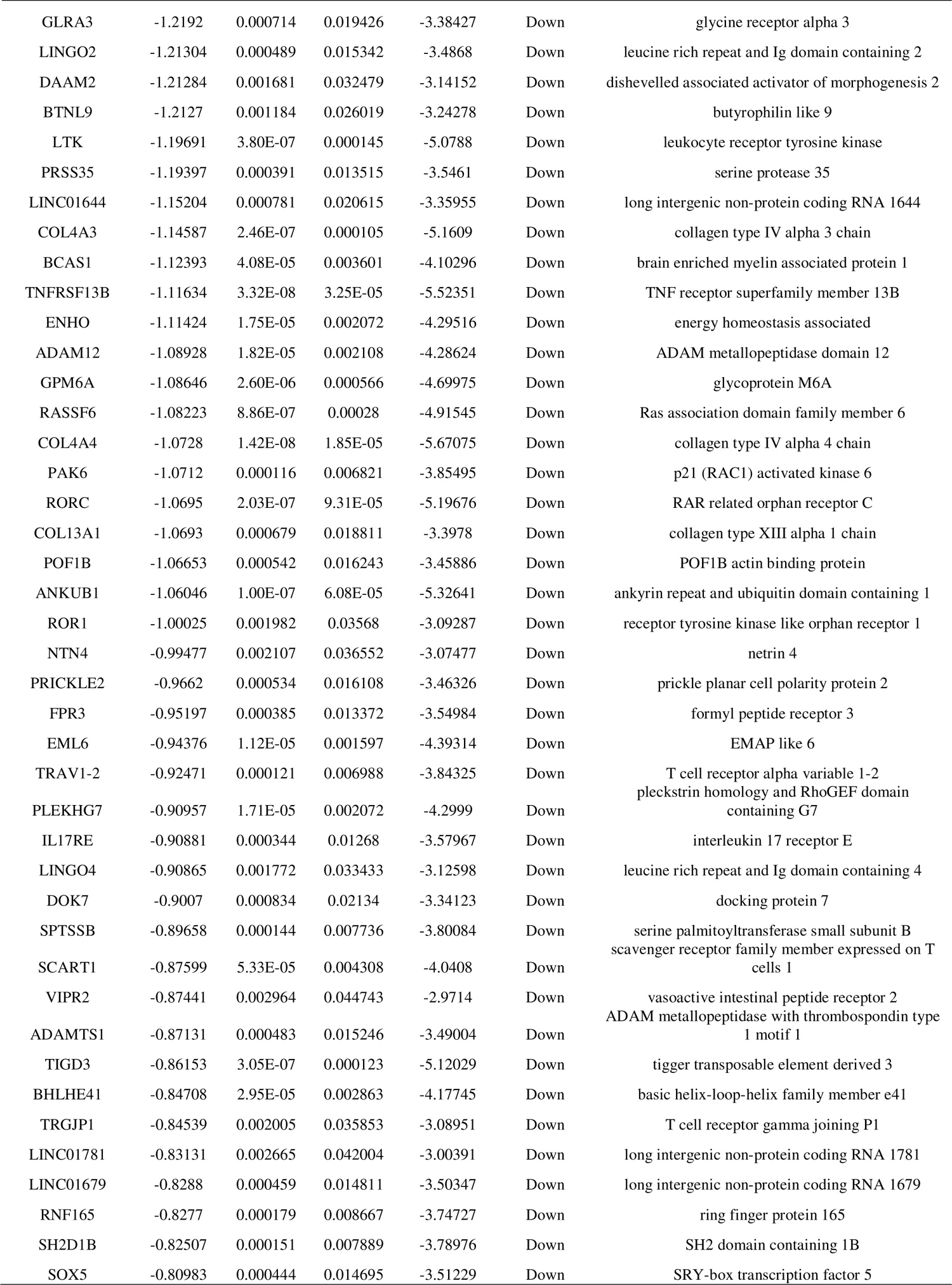
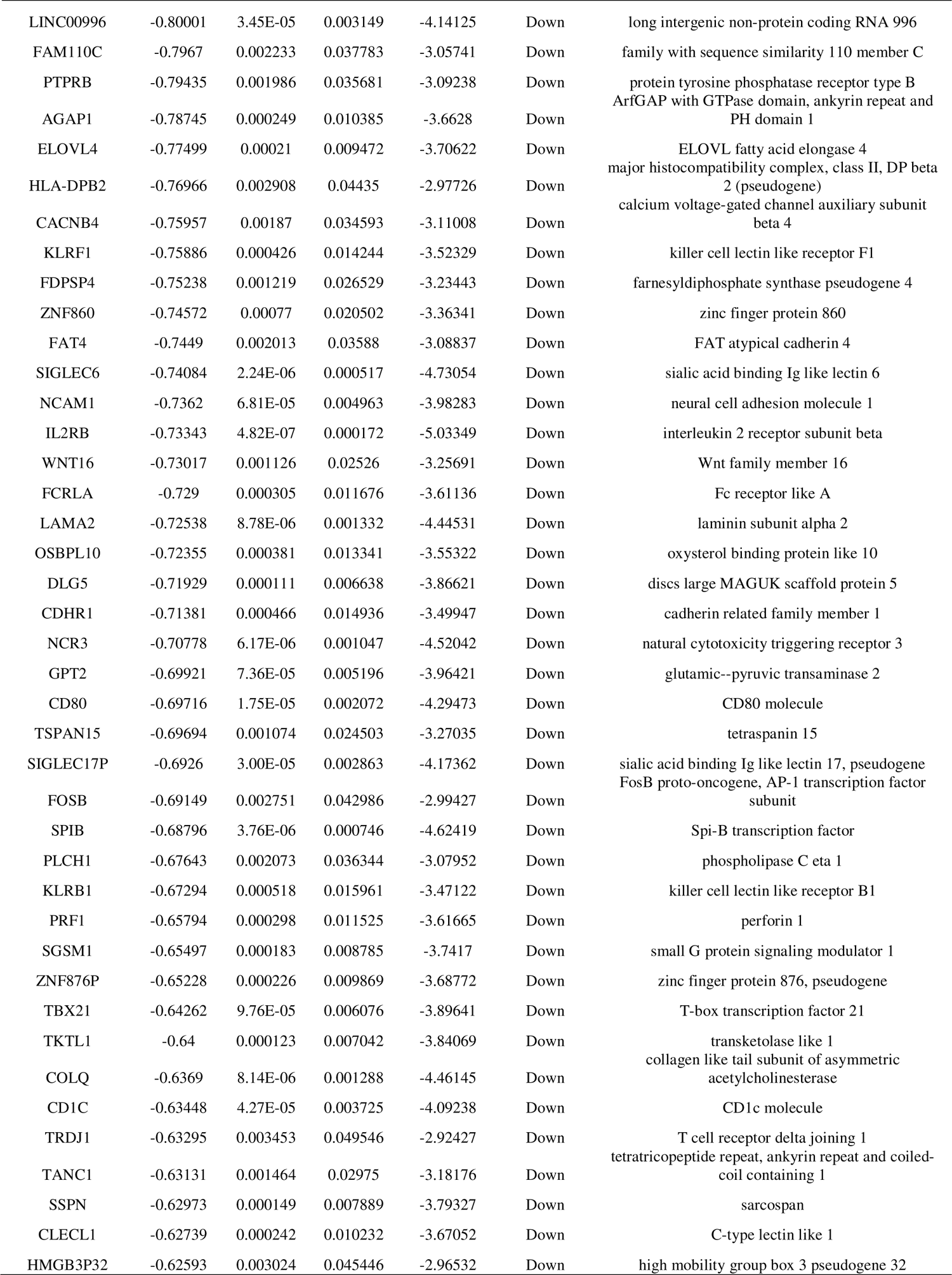
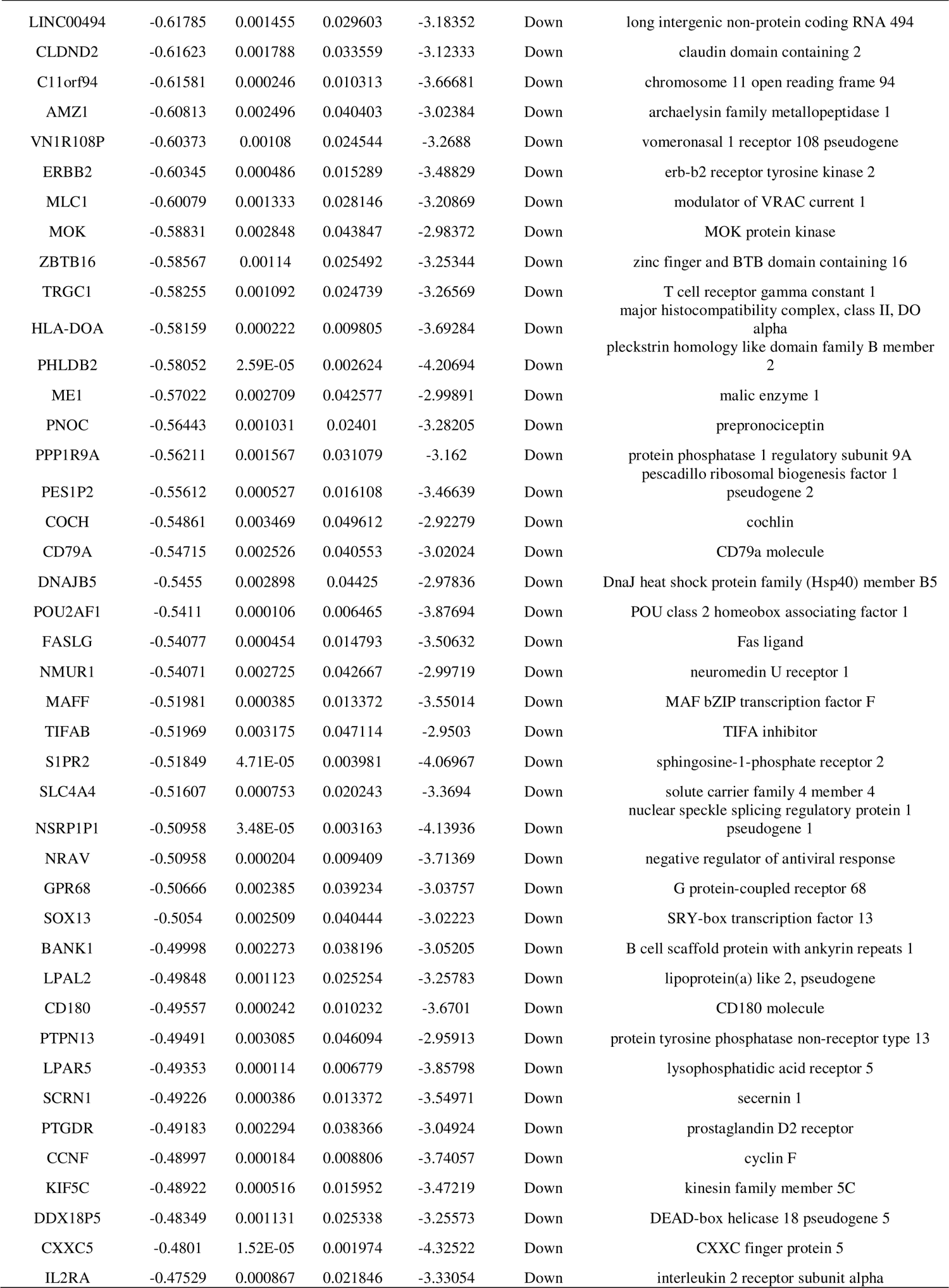
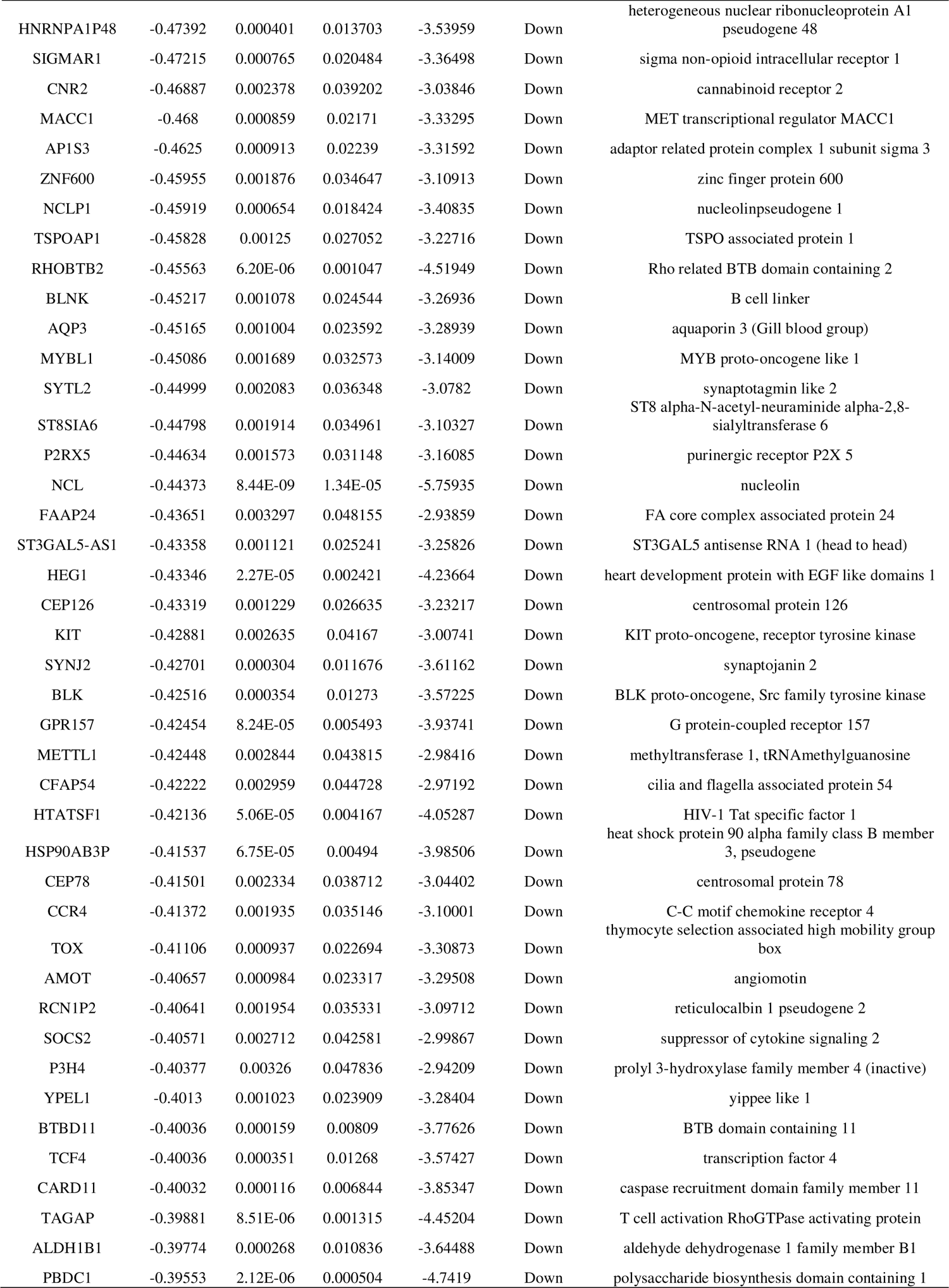
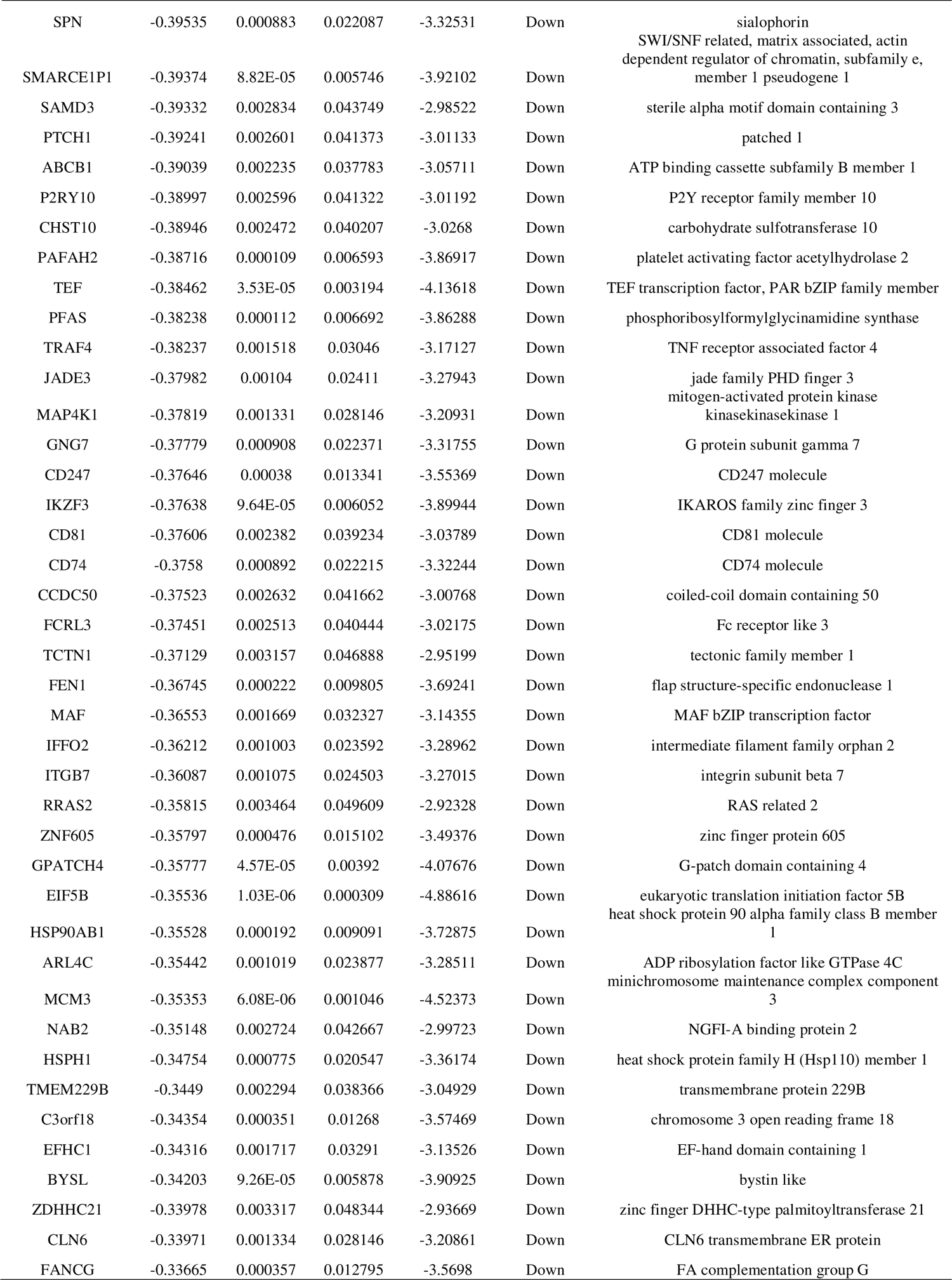
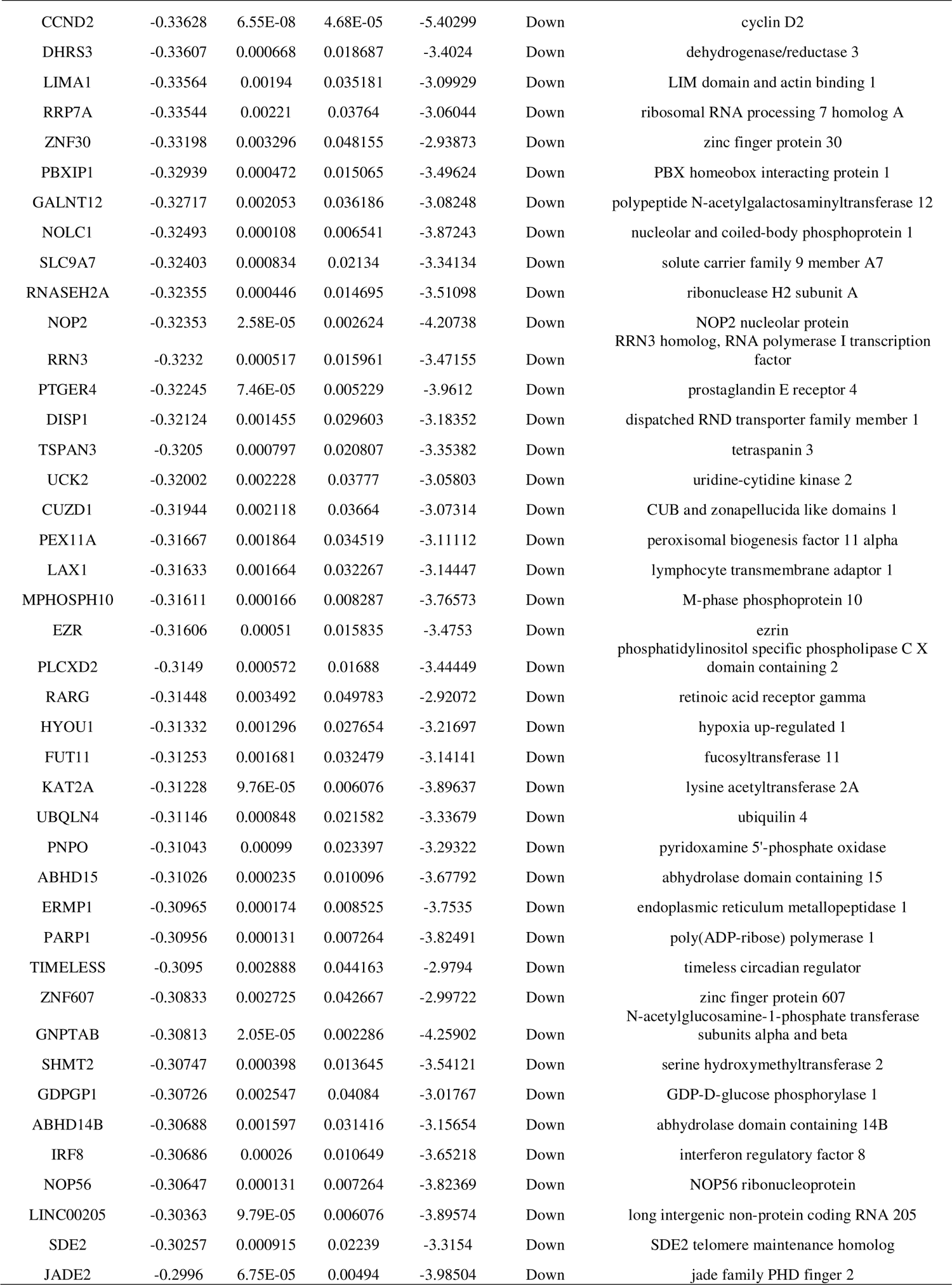
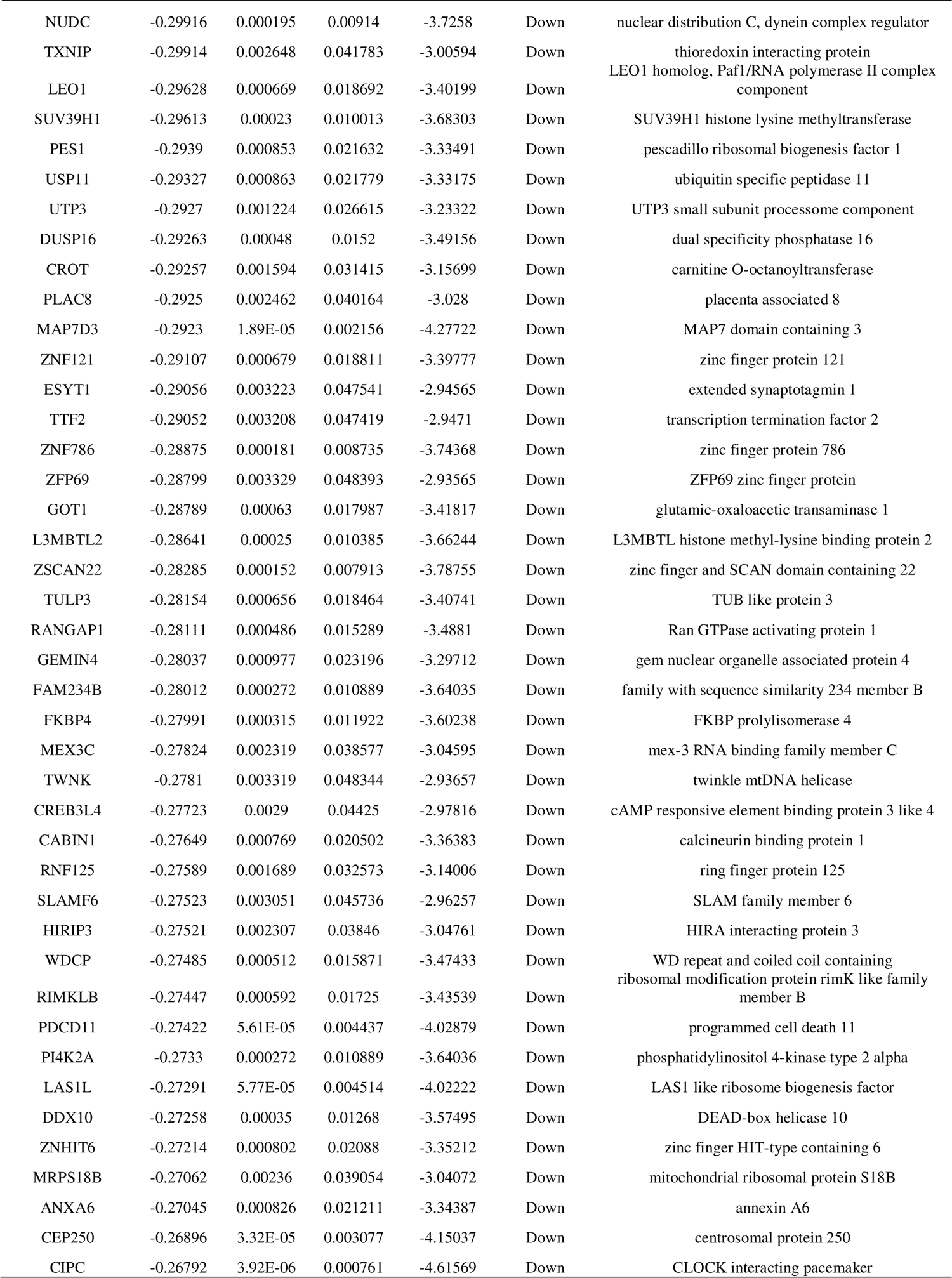
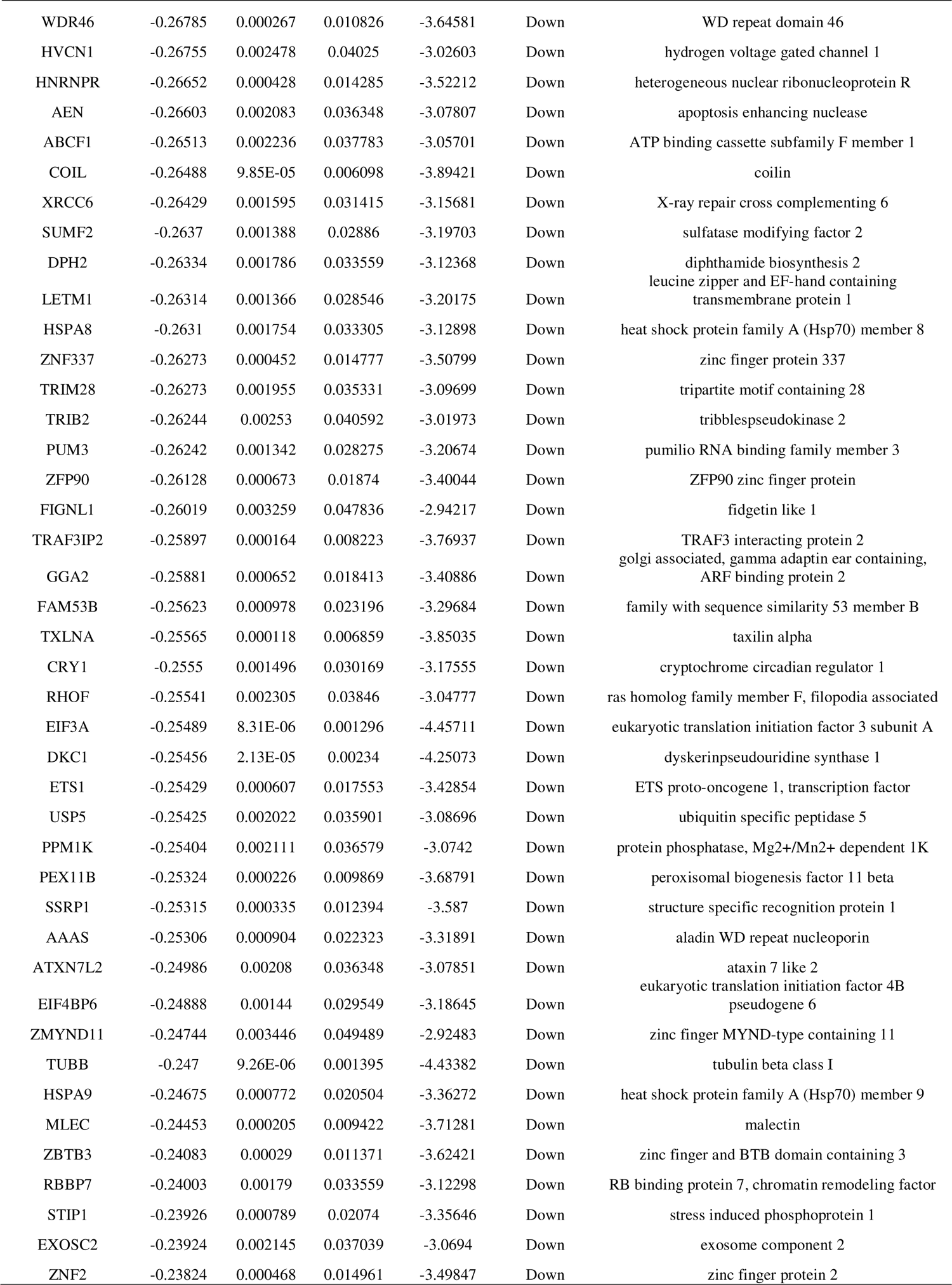
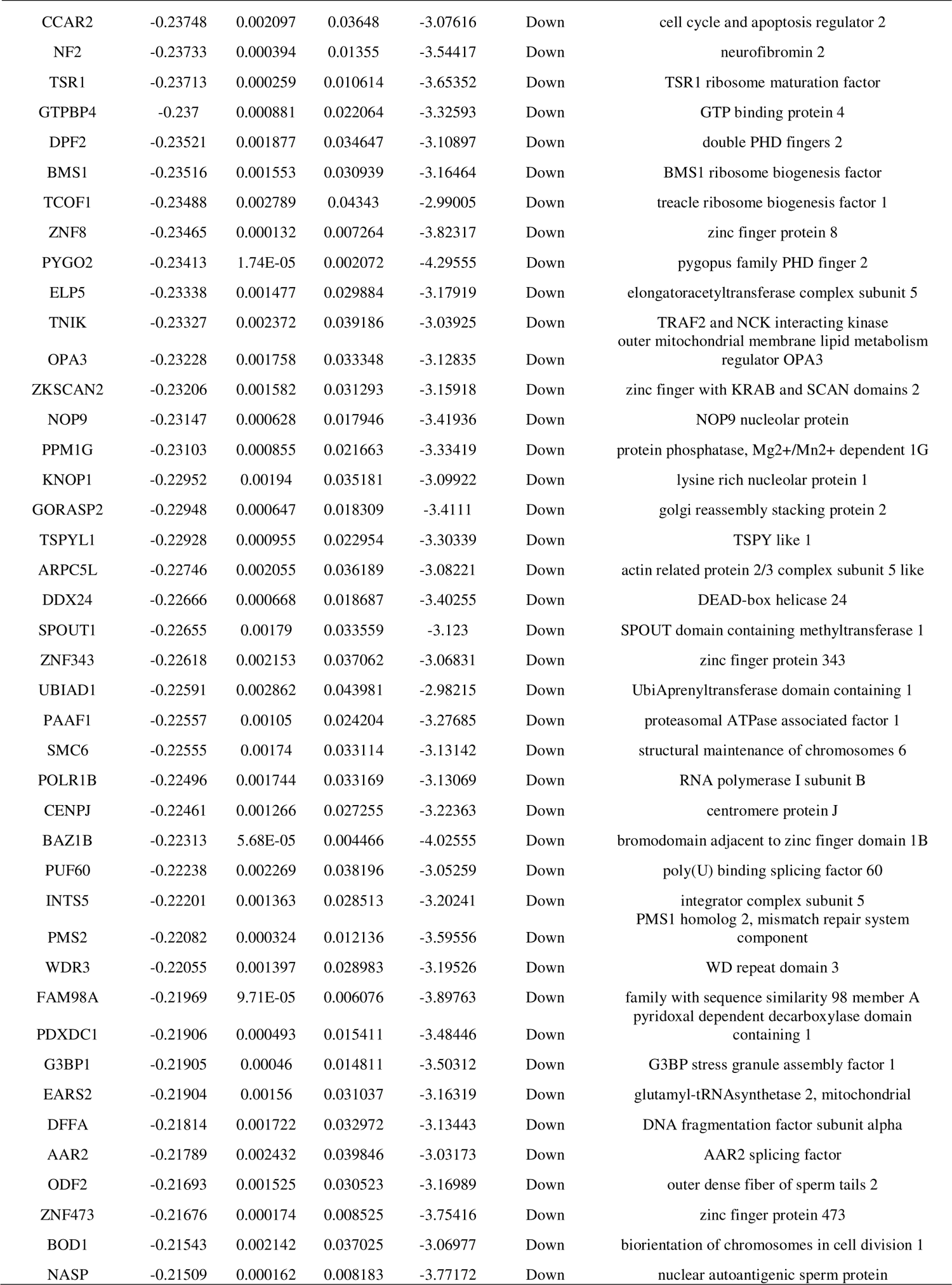
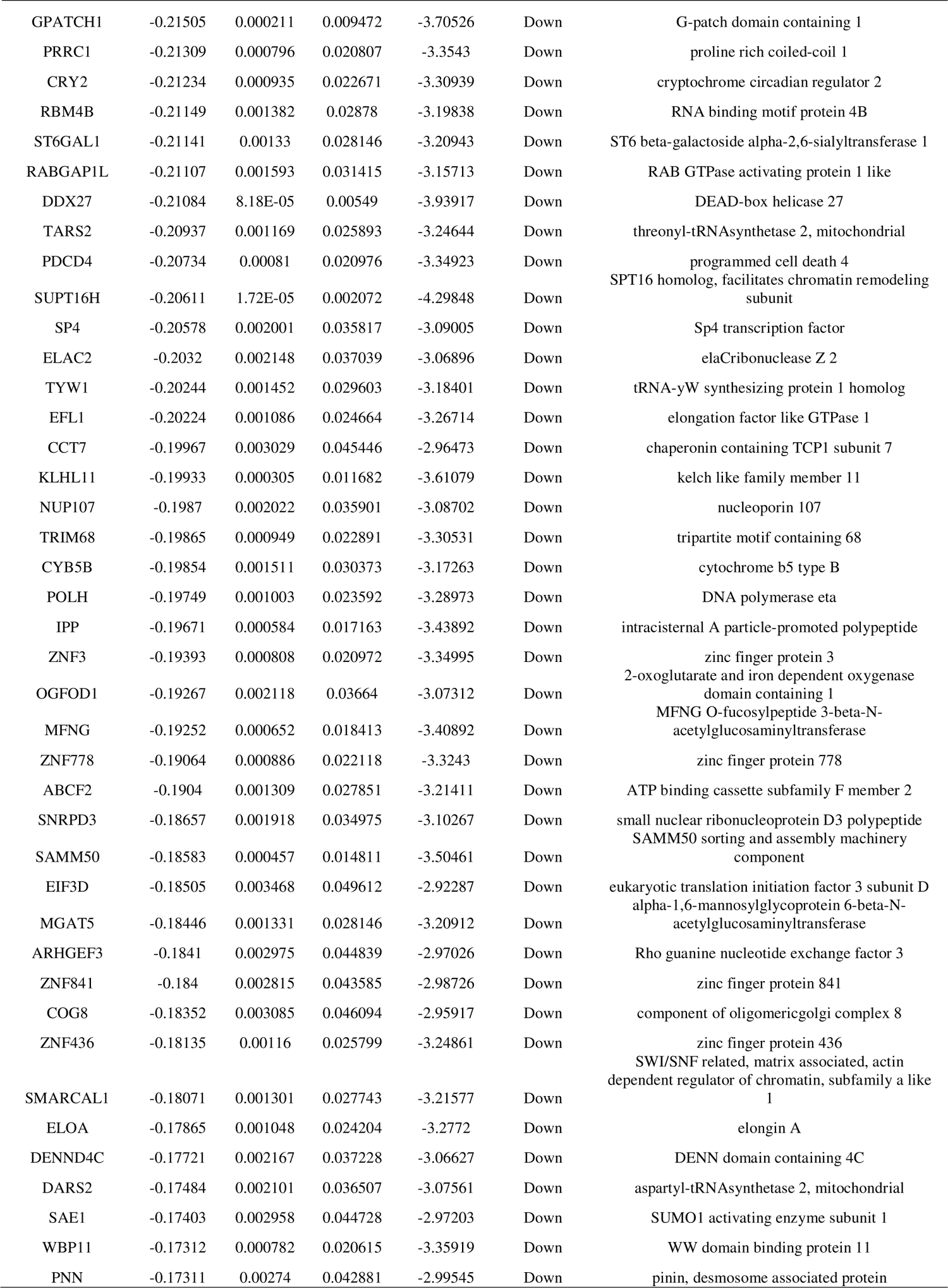
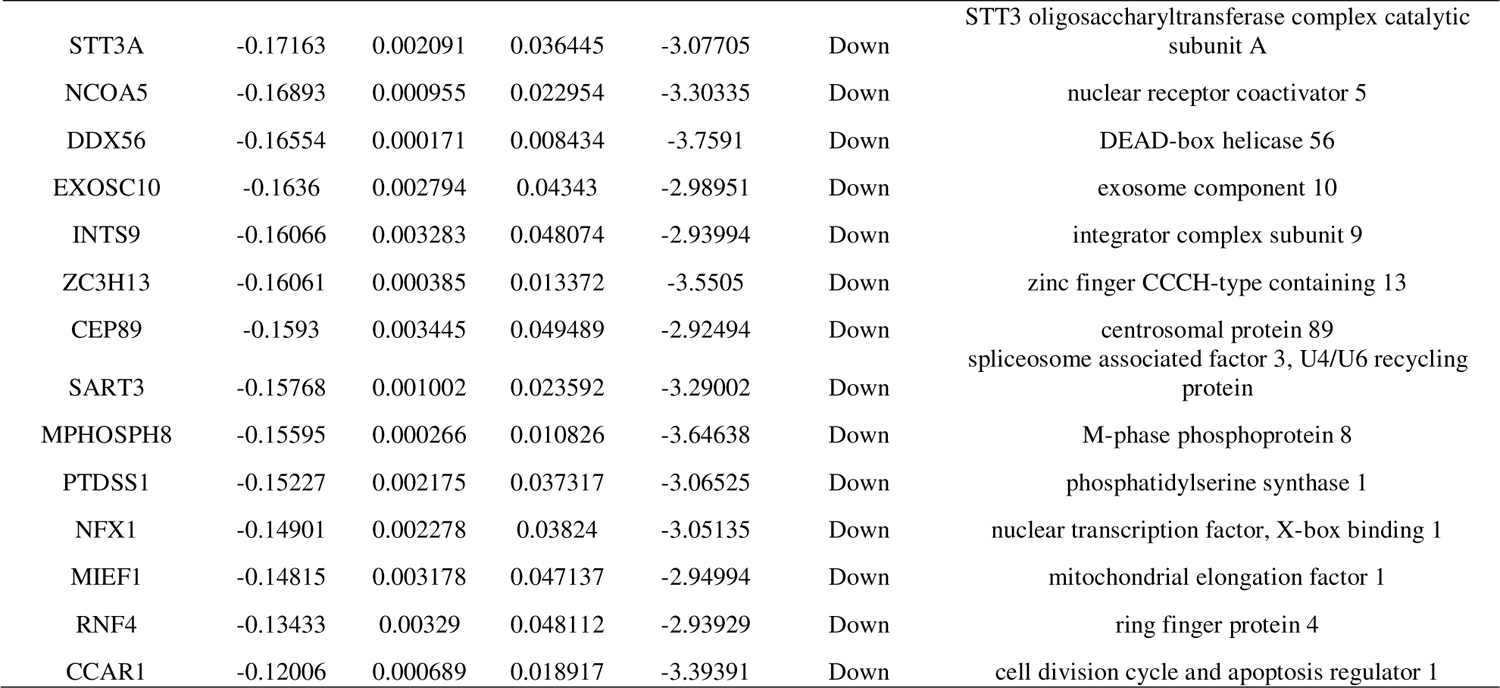
The statistical metrics for key differentially expressed genes (DEGs)

### GO and pathway enrichment analyses of DEGs

GO and REACTOME pathway enrichment analyses were performed using the g:Profiler website. The top terms of BP, CC, MF, and REACTOME pathways are shown in Table 2 and Table 3. As for BP, highly expressed DEGs were mainly enriched in response to stimulus and developmental process, whereas DEGs with low expression were mainly associated with regulation of cellular process and biological regulation. In the analysis of the CC enrichment, highly expressed DEGs correlated with plasma membrane and cytoplasm. For another, lowly expressed DEGs were predominant in nucleus and membrane-bounded organelle. About MF, highly expressed DEGs were primarily enriched in catalytic activity and ion binding, whereas DEGs with low expression were mostly enriched in organic cyclic compound binding and nucleic acid binding. However, for REACTOME pathway enrichment, highly expressed DEGs mainly involved in neutrophil degranulation and immune system, whereas lowly expressed DEGs mainly involved in rRNA processing and metabolism of RNA.

**Table 2.**
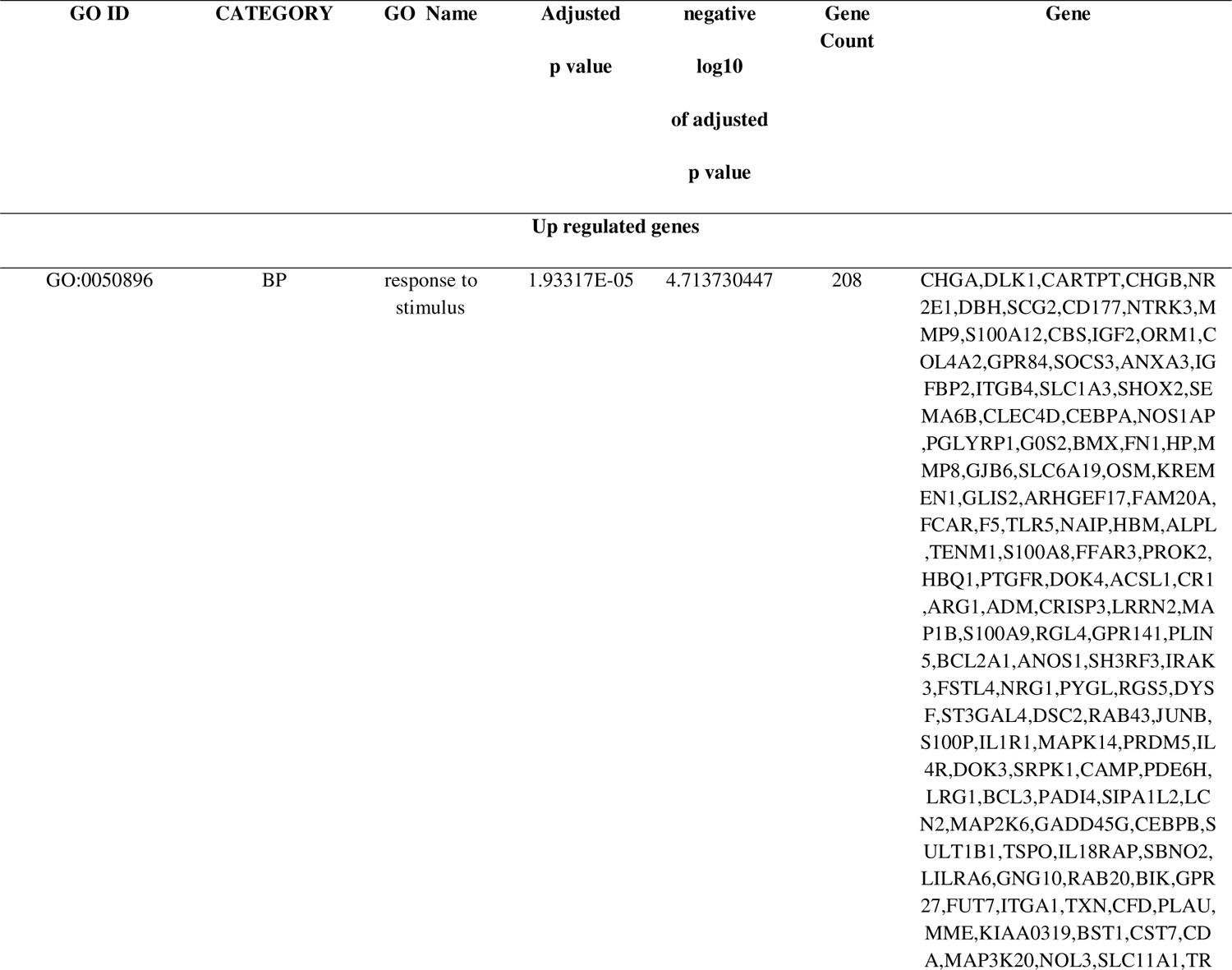
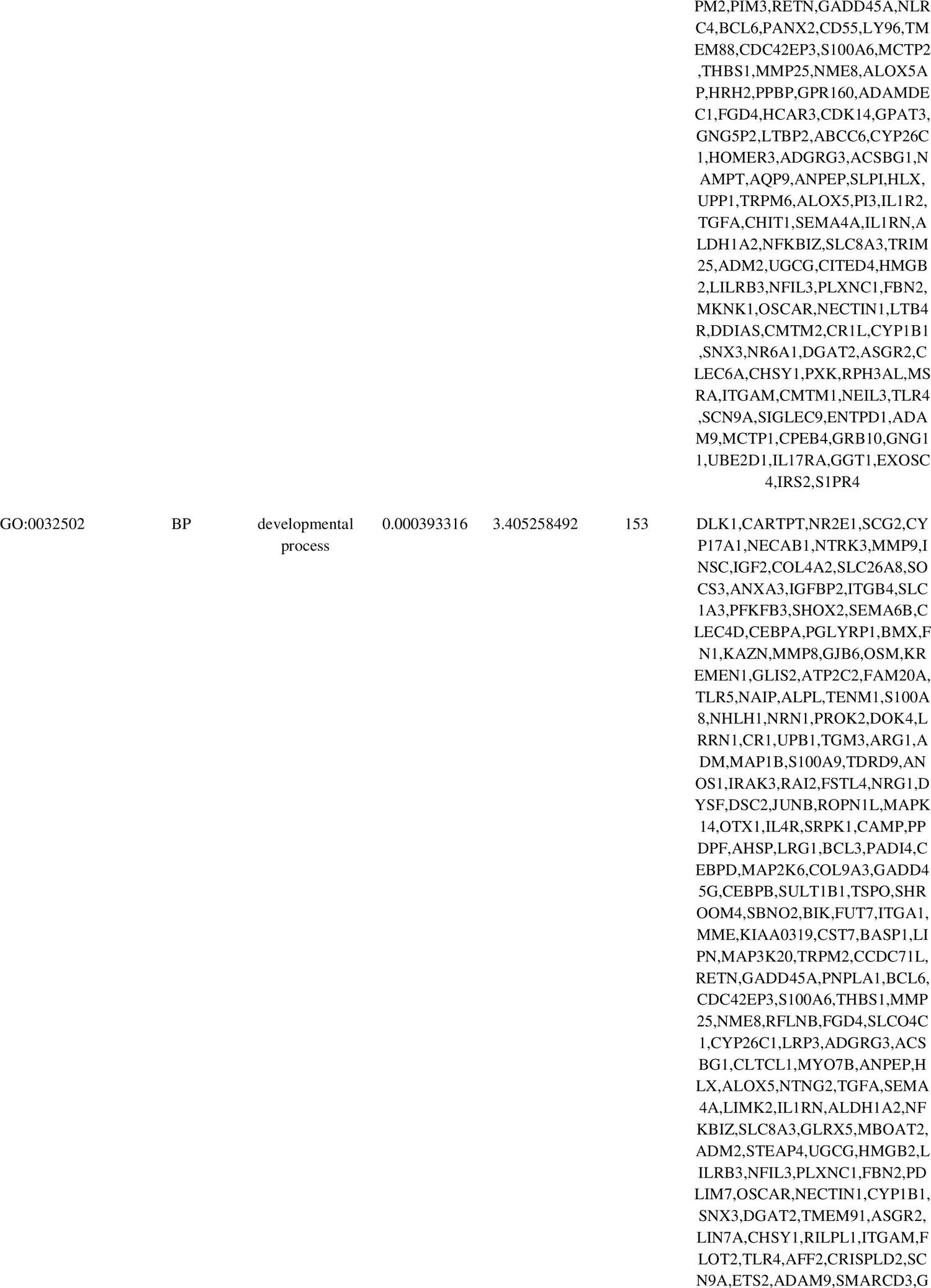
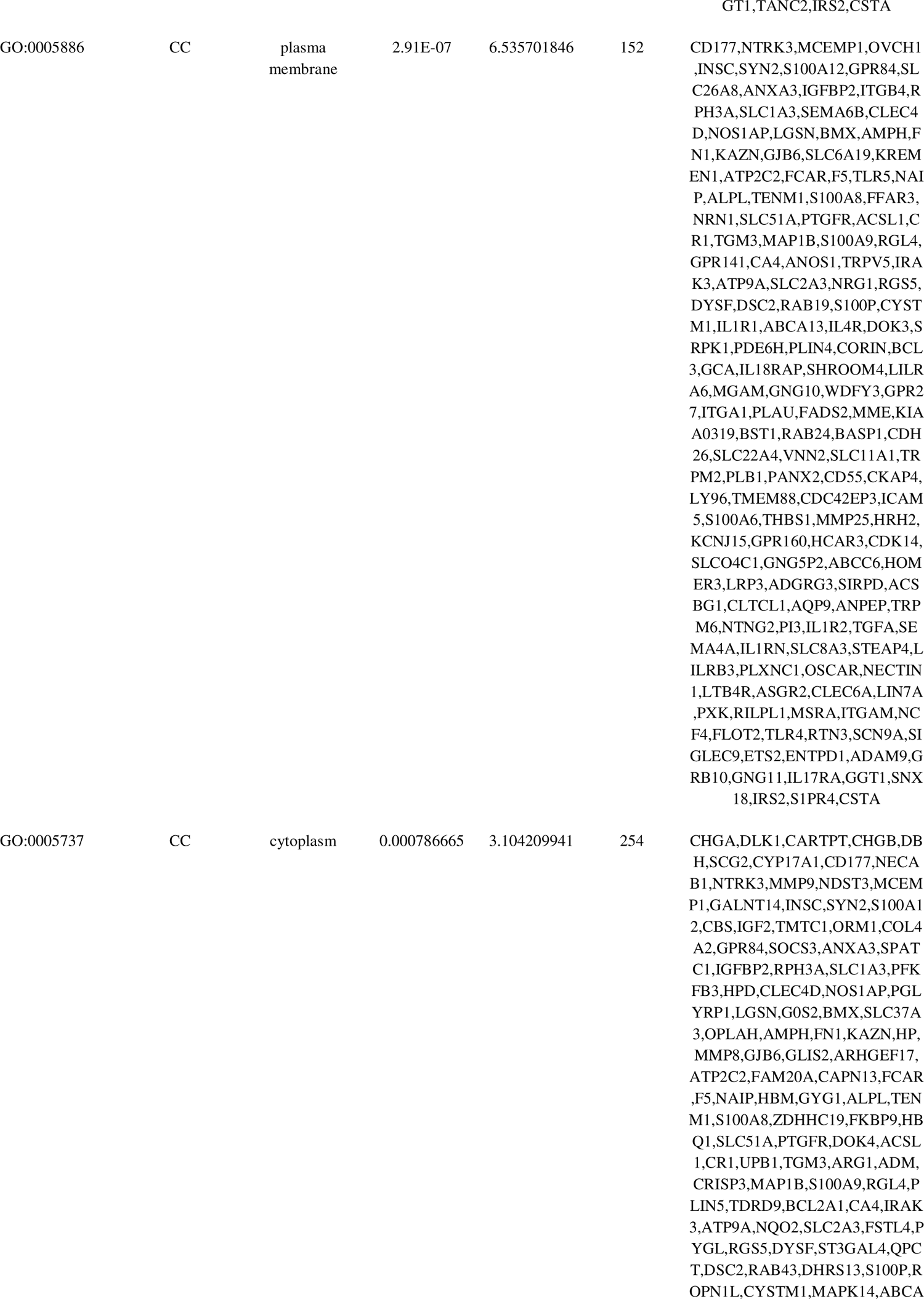
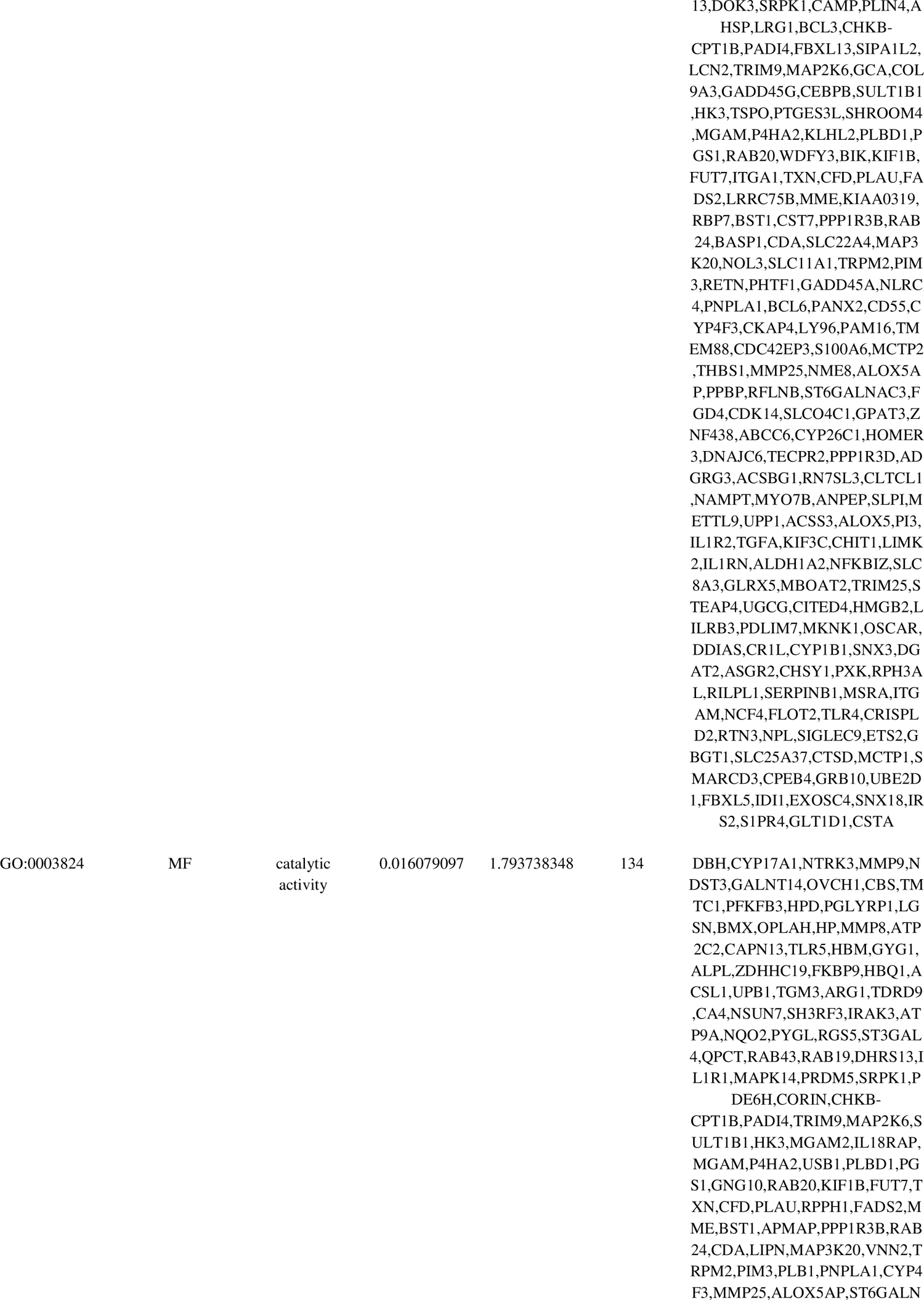
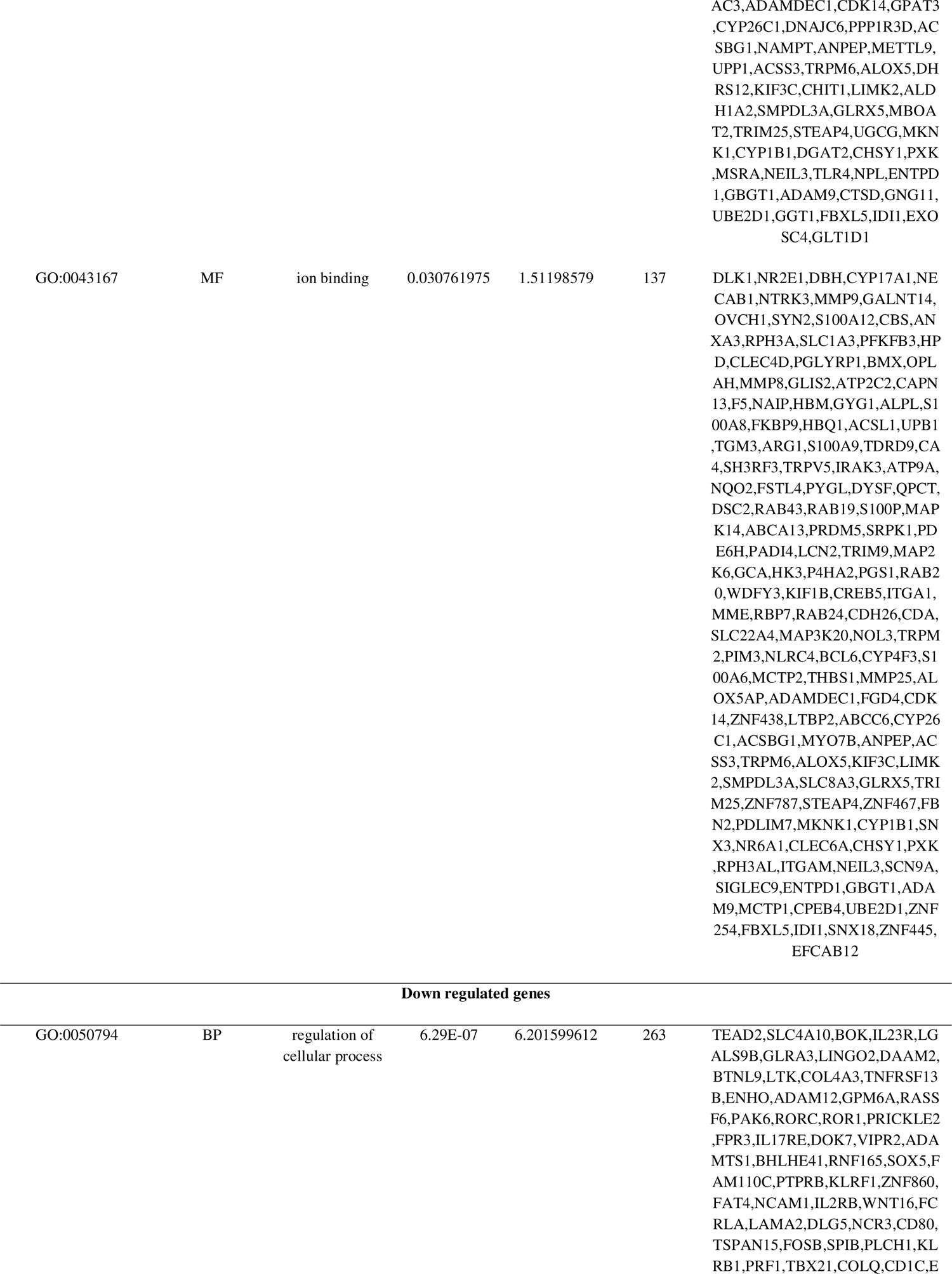
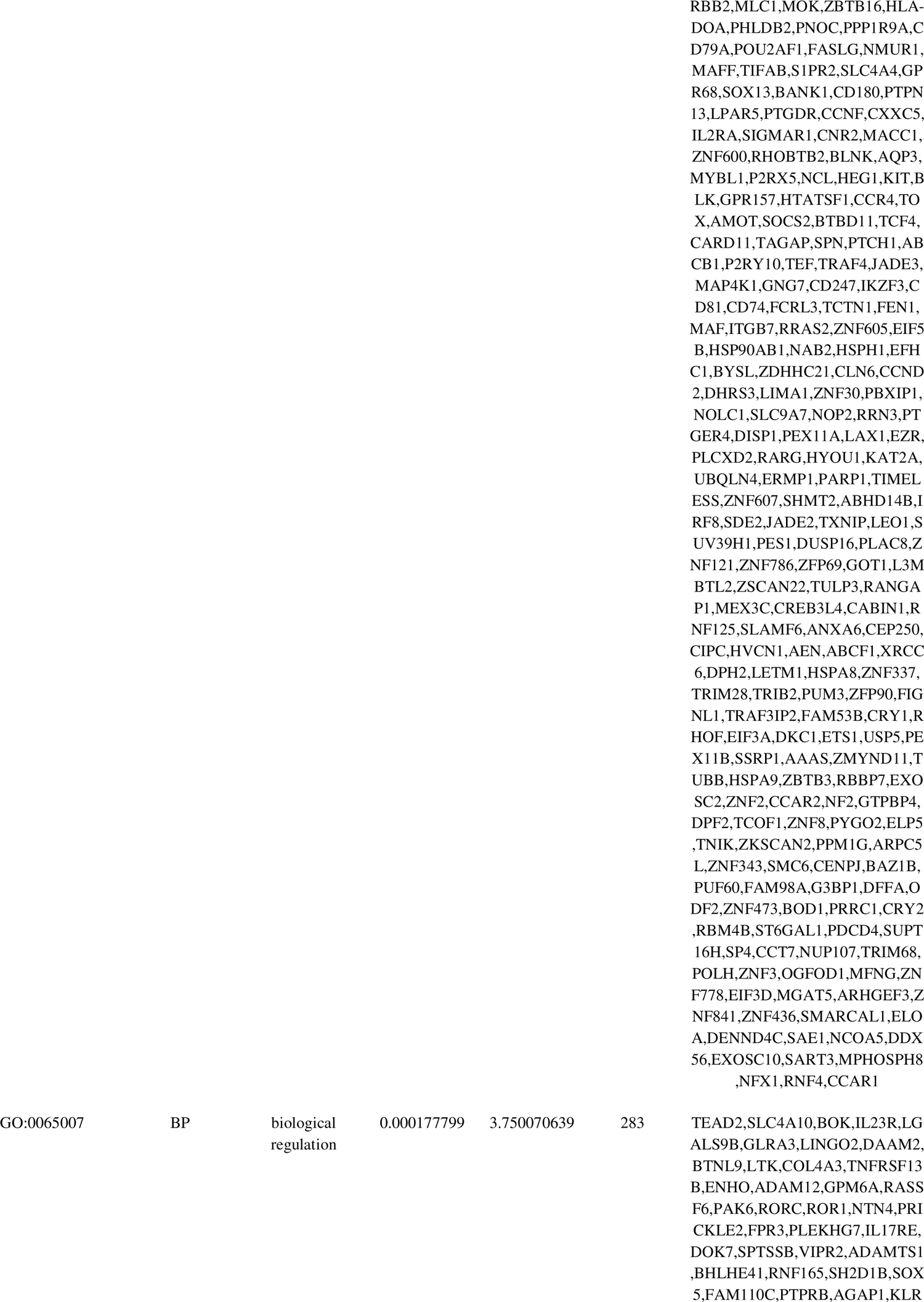
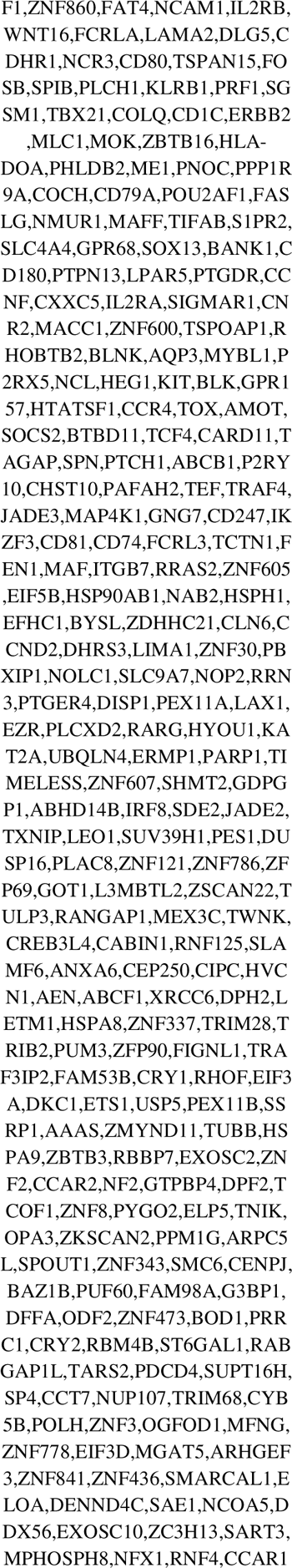
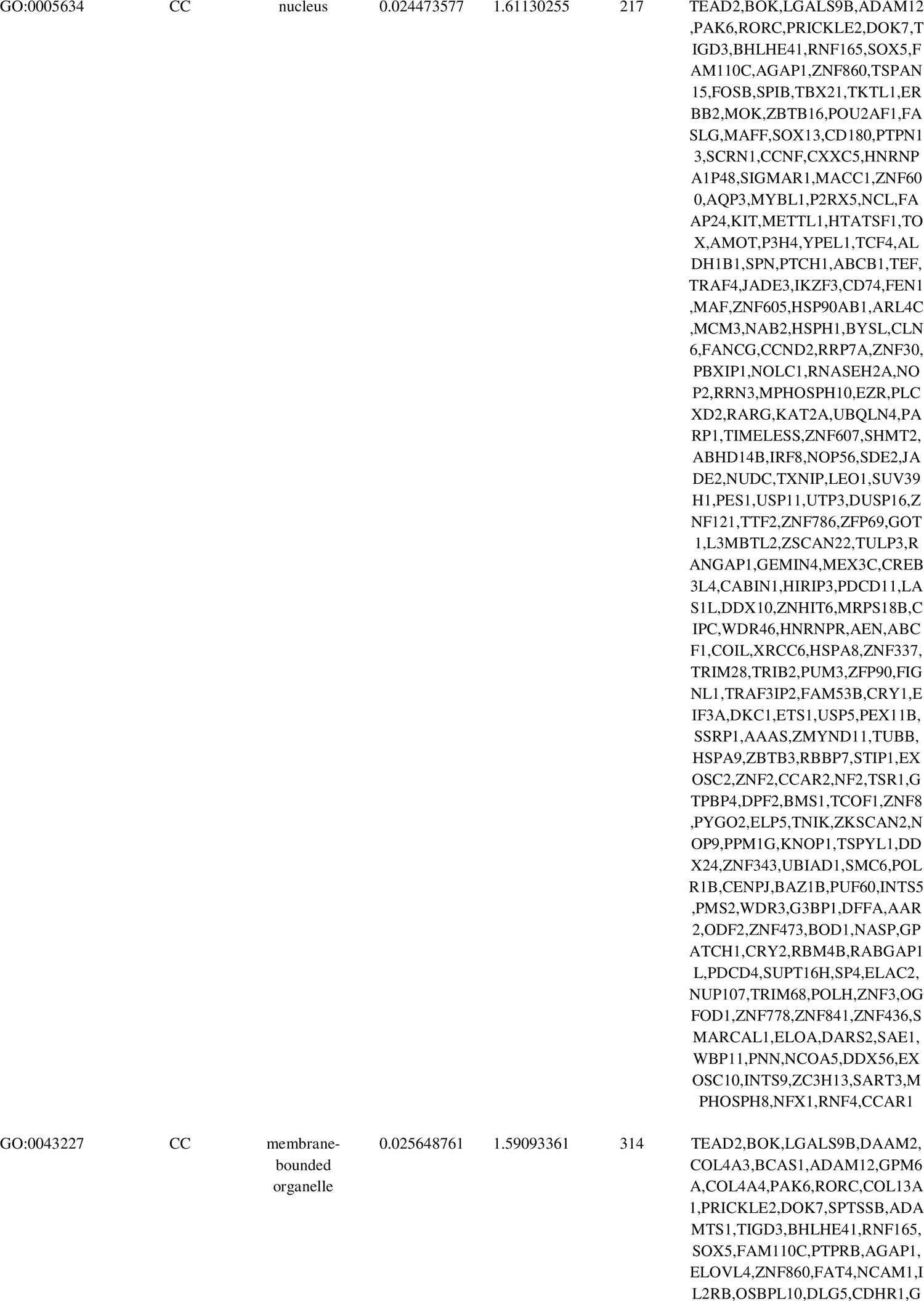
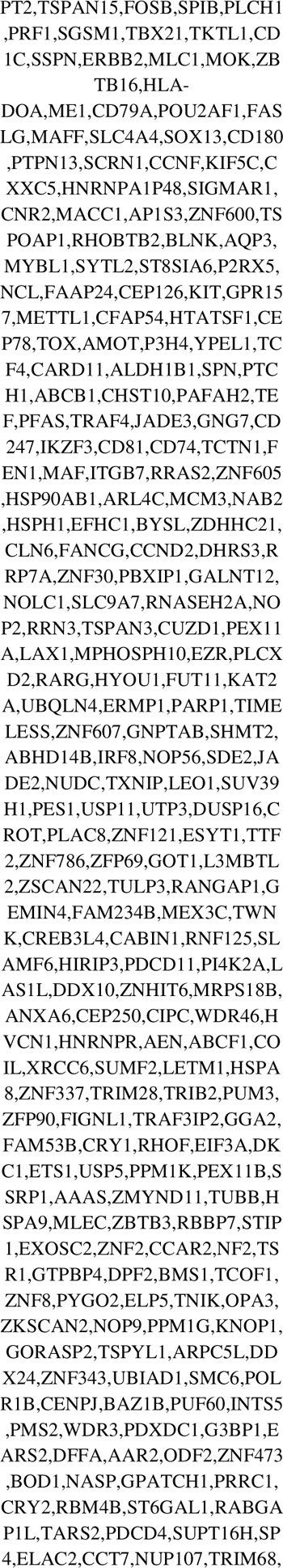
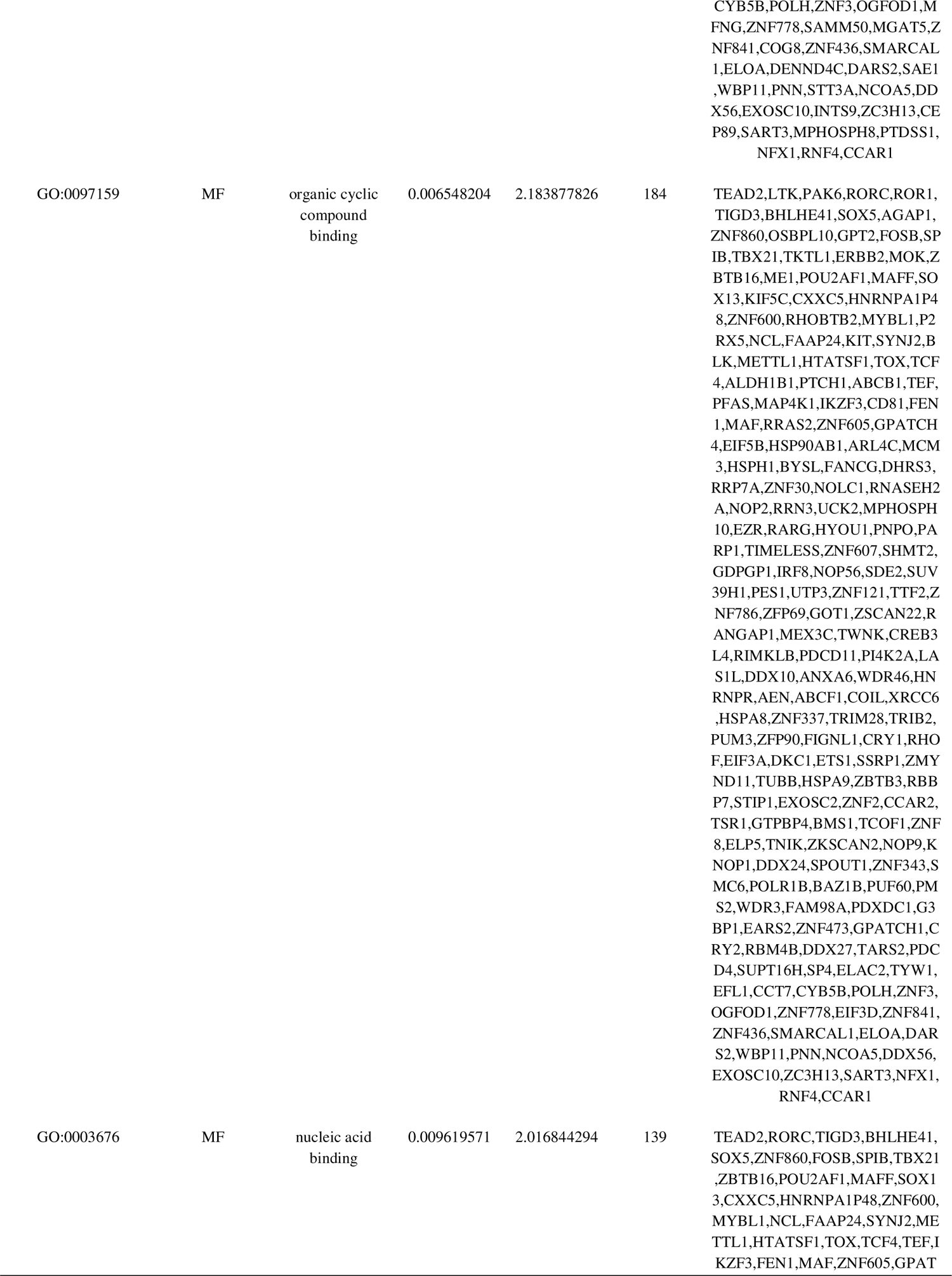
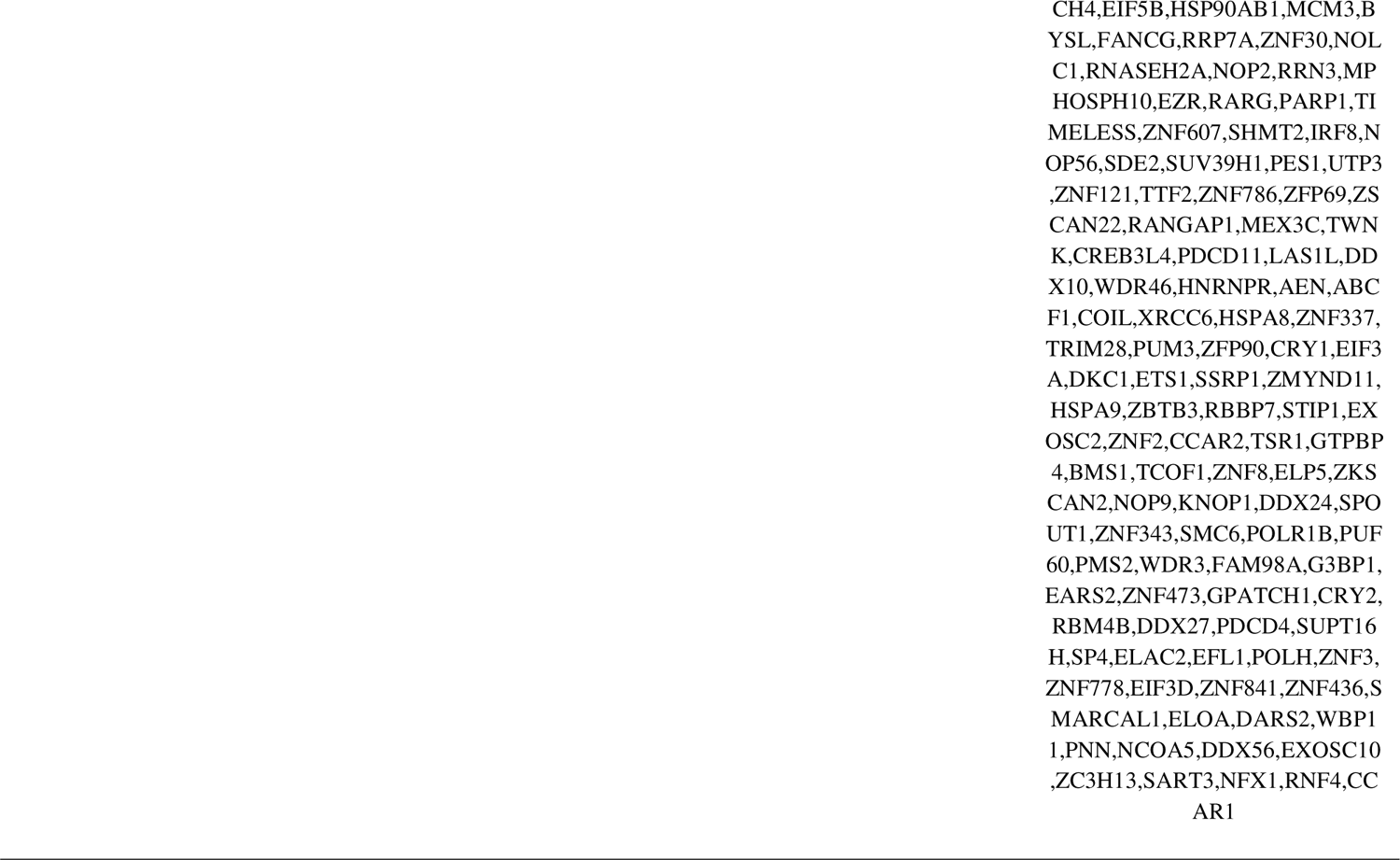
The enriched GO terms of the up and down regulated differentially expressed genes

**Table 3.**
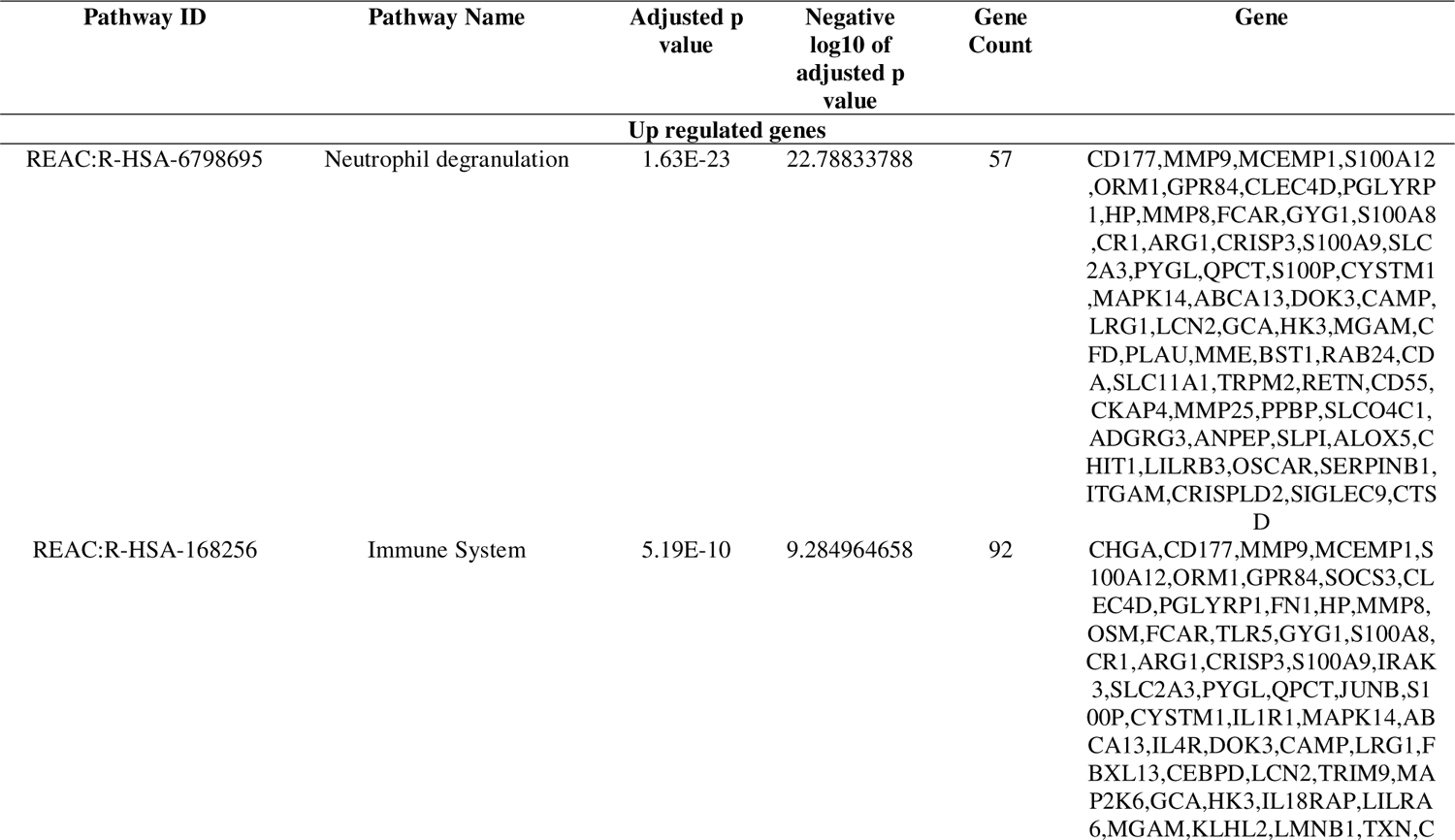
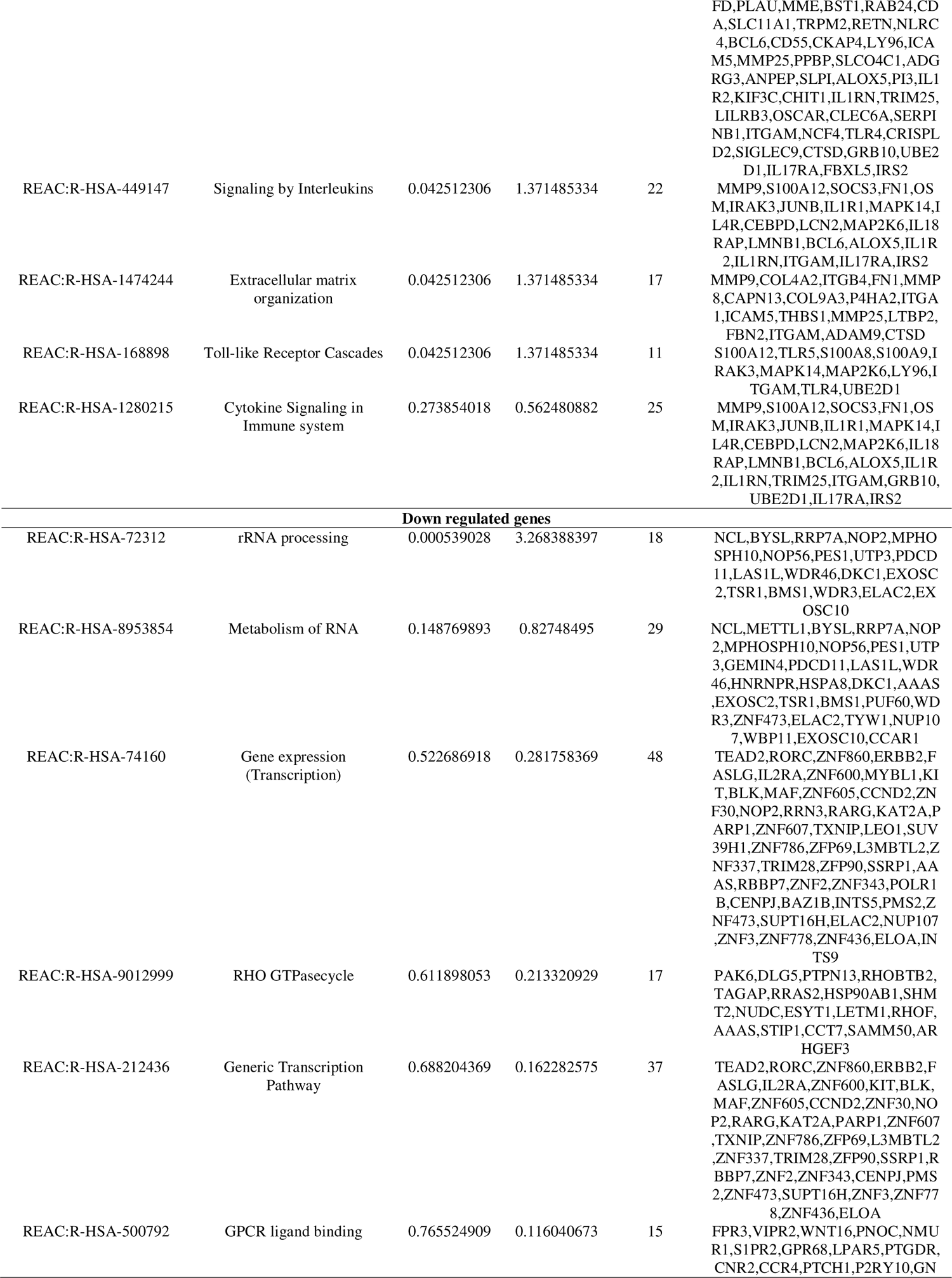
The enriched pathway terms of the up and down regulated differentially expressed genes

### Construction of the PPI network and module analysis

Based on the results of IMEx interactome online tool analysis of the DEGs, a PPI network was constructed by the Cytoscape software, which comprised 4993 nodes and 11778 edges (Fig. 3). DEGs with high node degree, betweenness, stress and closeness were selected as the hub genes in CF from the PPI network: FN1, UBE2D1, SRPK1, MAPK14, CEBPB, HSP90AB1, HSPA8, XRCC6, NCL and PARP1. The score of each hub gene was shown in Table 4. The PEWCC1 plug-in in Cytoscape is used to construct functional modules. A total of 53 nodes and 127 edges were involved in module 1. The most significant module 1 of up regulated genes is shown in Fig.4A, and it was mainly linked to response to stimulus, immune system, developmental process, signaling by interleukins and cytoplasm. A total of 47 nodes and 130 edges were involved in module 2. The most significant module 2 of down regulated genes is shown in Fig.4B, and it was mainly linked to rRNA processing, metabolism of RNA and regulation of cellular process.

**Fig. 3.**
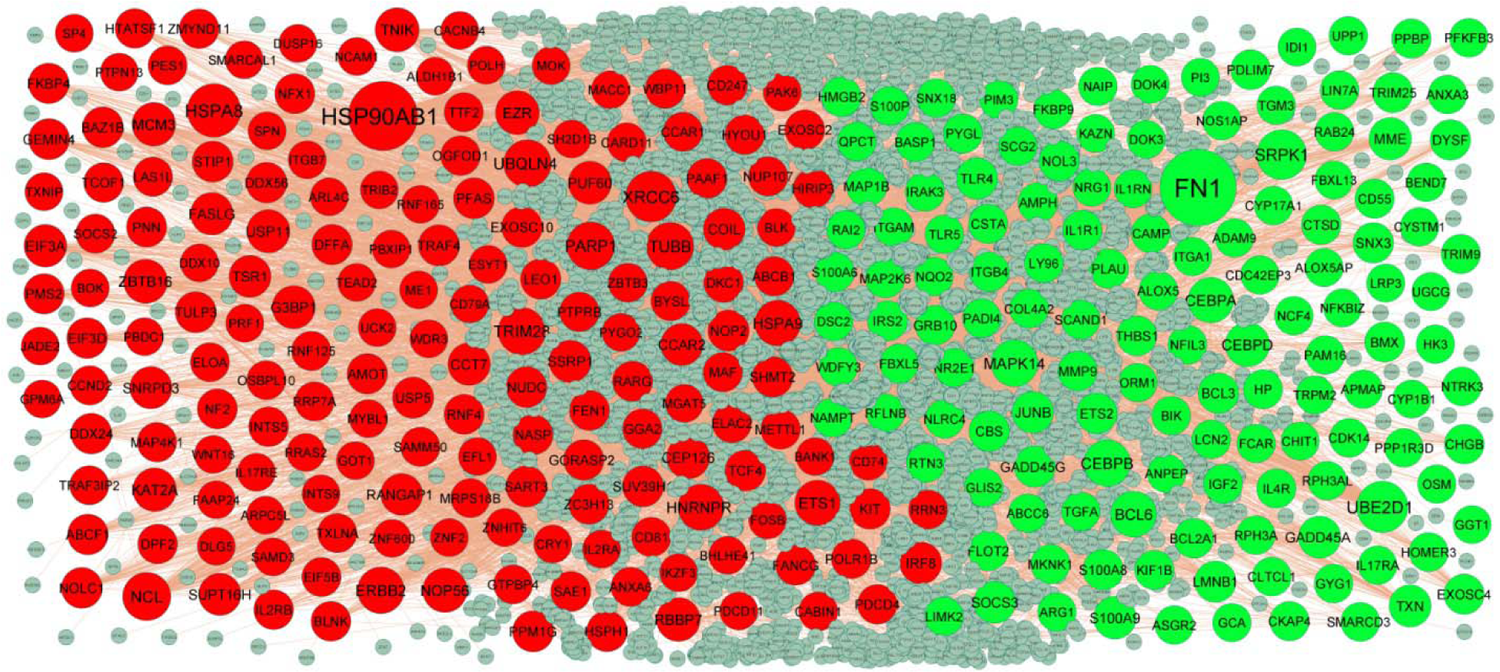
PPI network of DEGs. Up regulated genes are marked in green; down regulated genes are marked in red

**Fig. 4.**
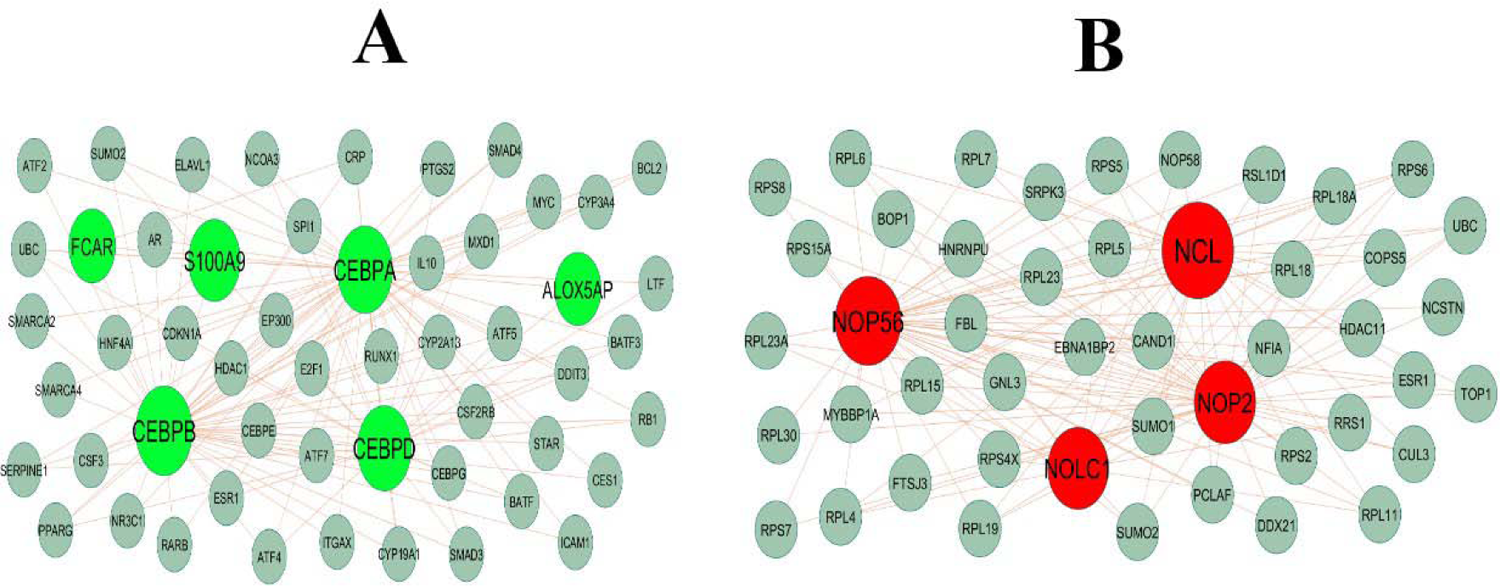
Modules selected from the DEG PPI between patients with FSGS and normal controls. (A) The most significant module was obtained from PPI network with 53 nodes and 127 edges for up regulated genes (B) The most significant module was obtained from PPI network with 47 nodes and 130 edges for down regulated genes. Up regulated genes are marked in green; down regulated genes are marked in red

**Table 4.**
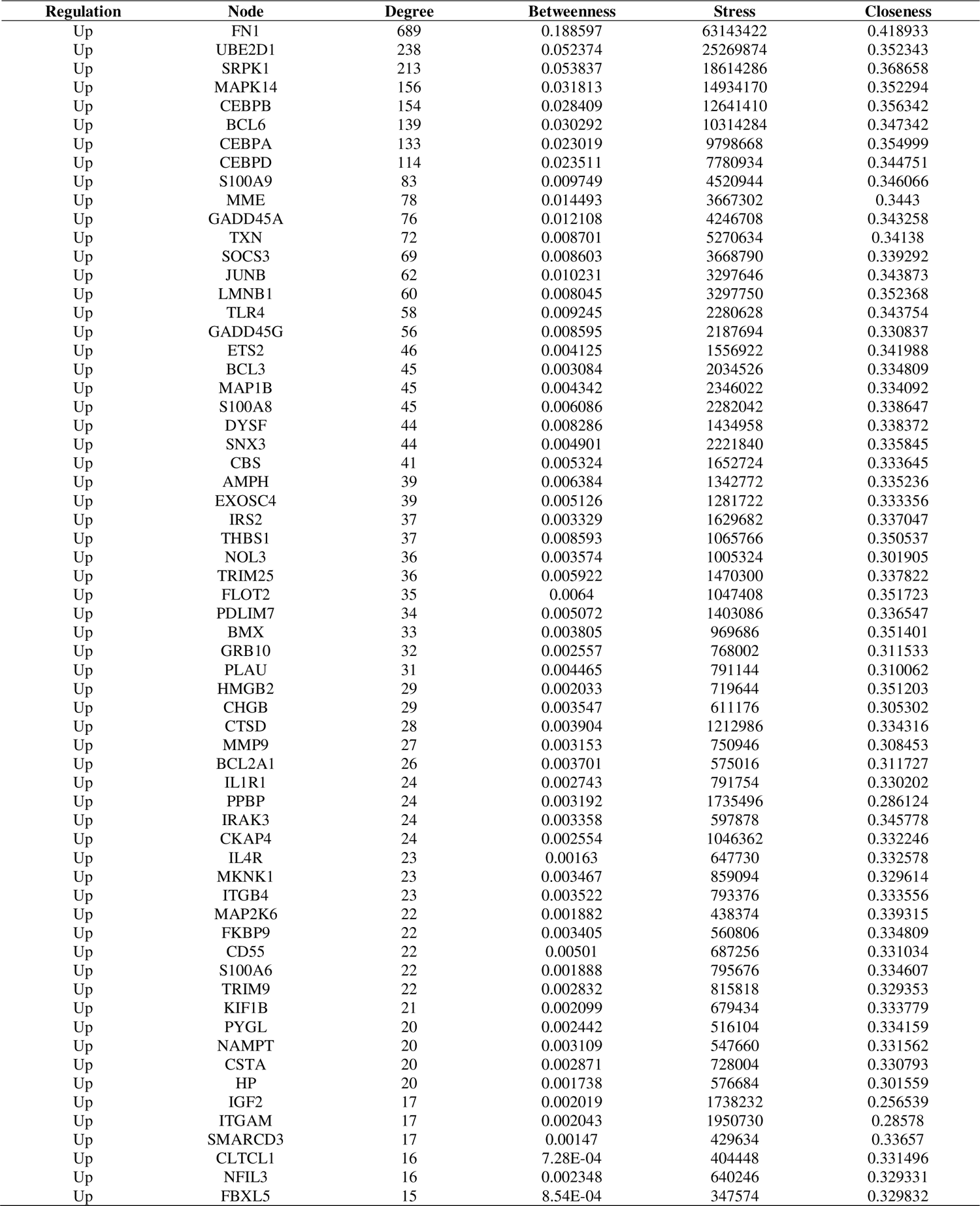
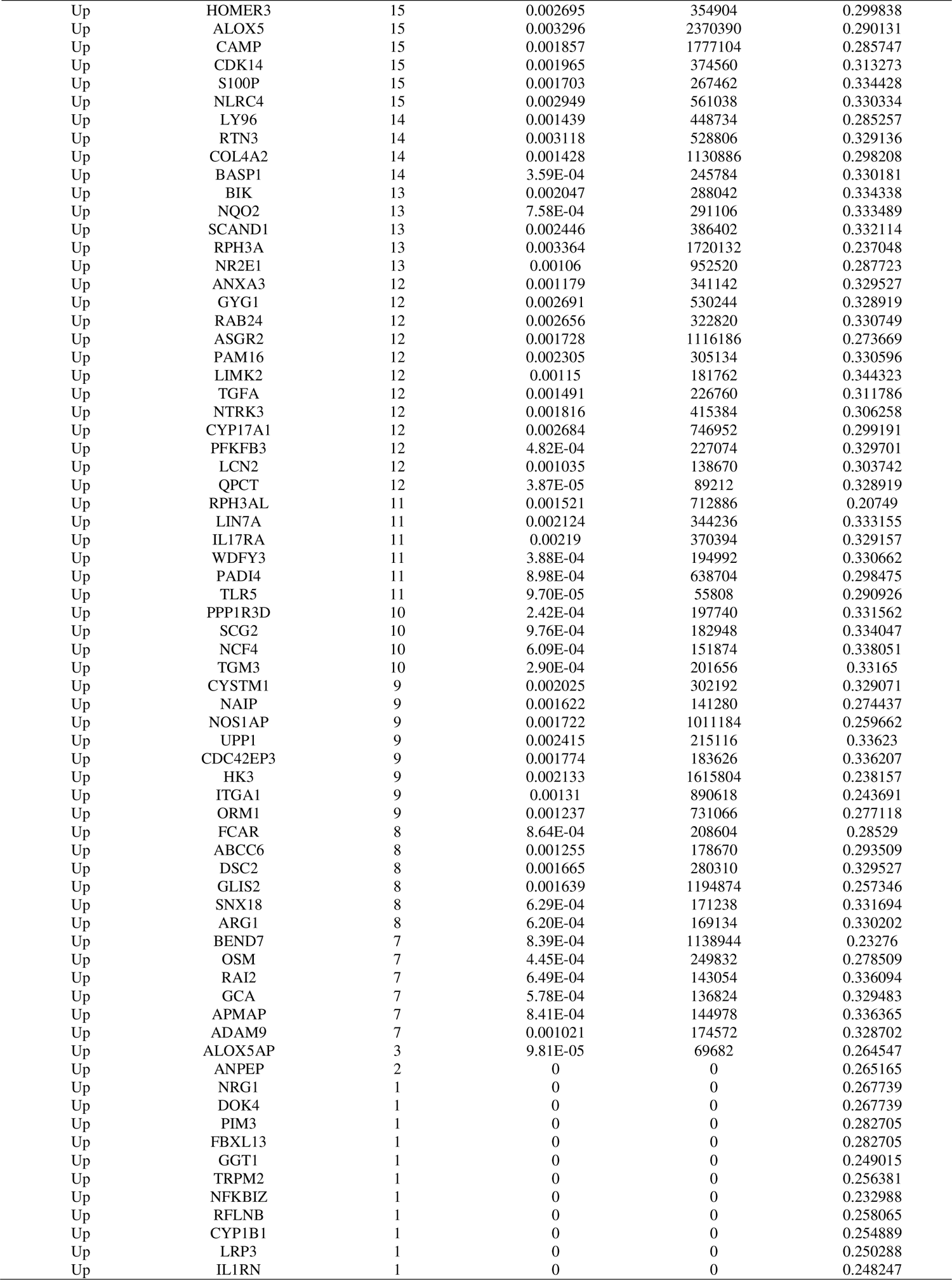
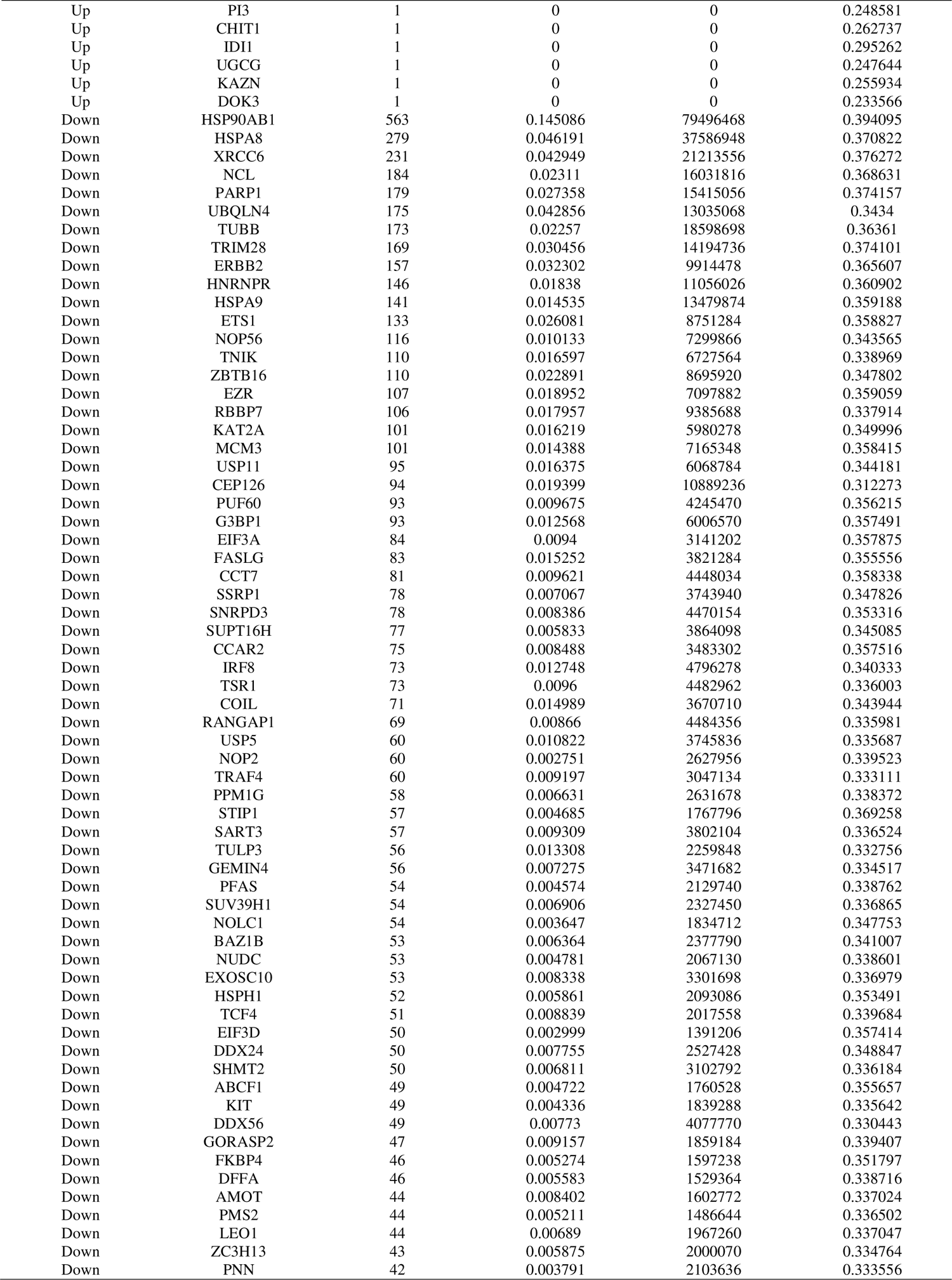
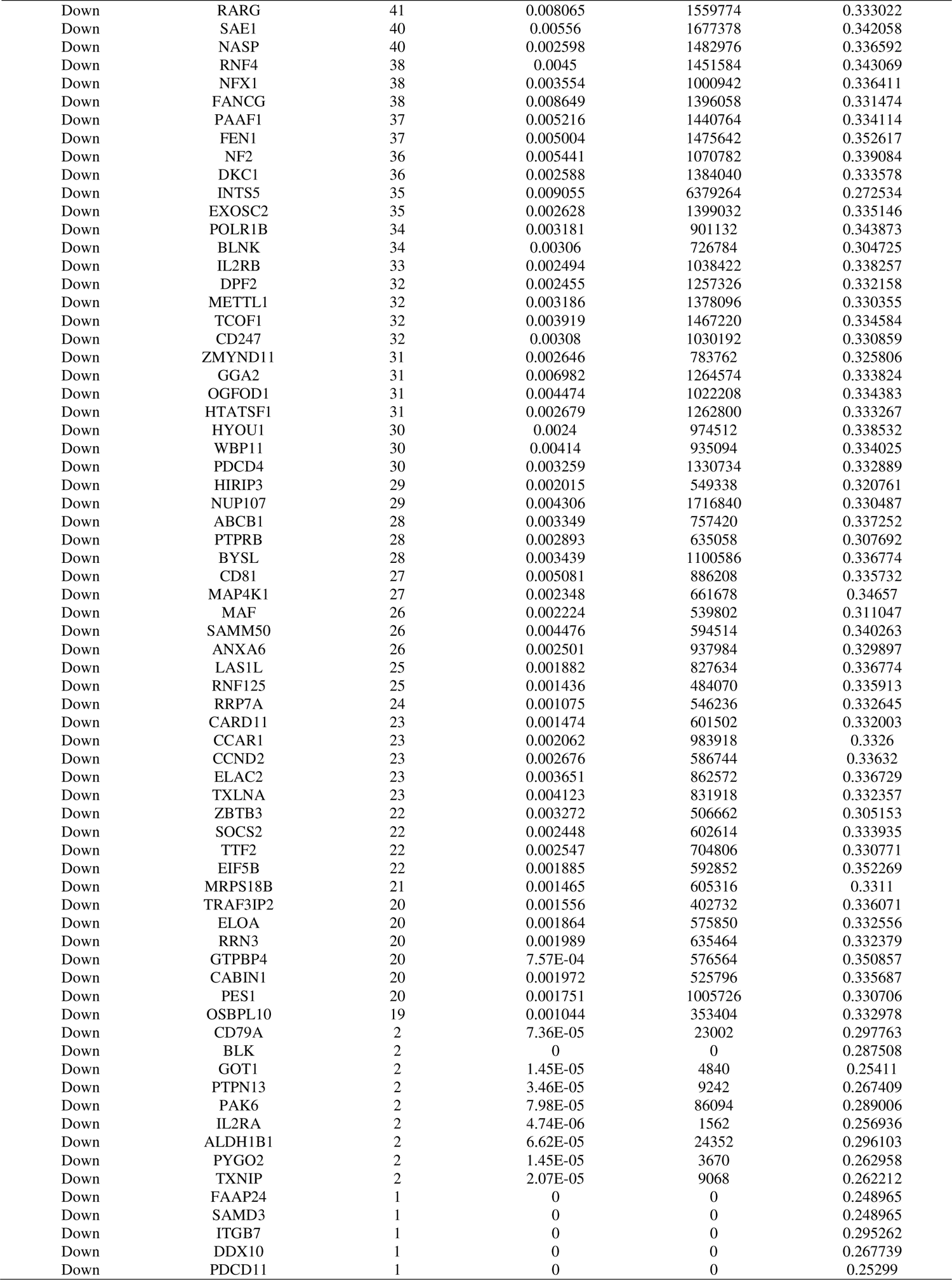
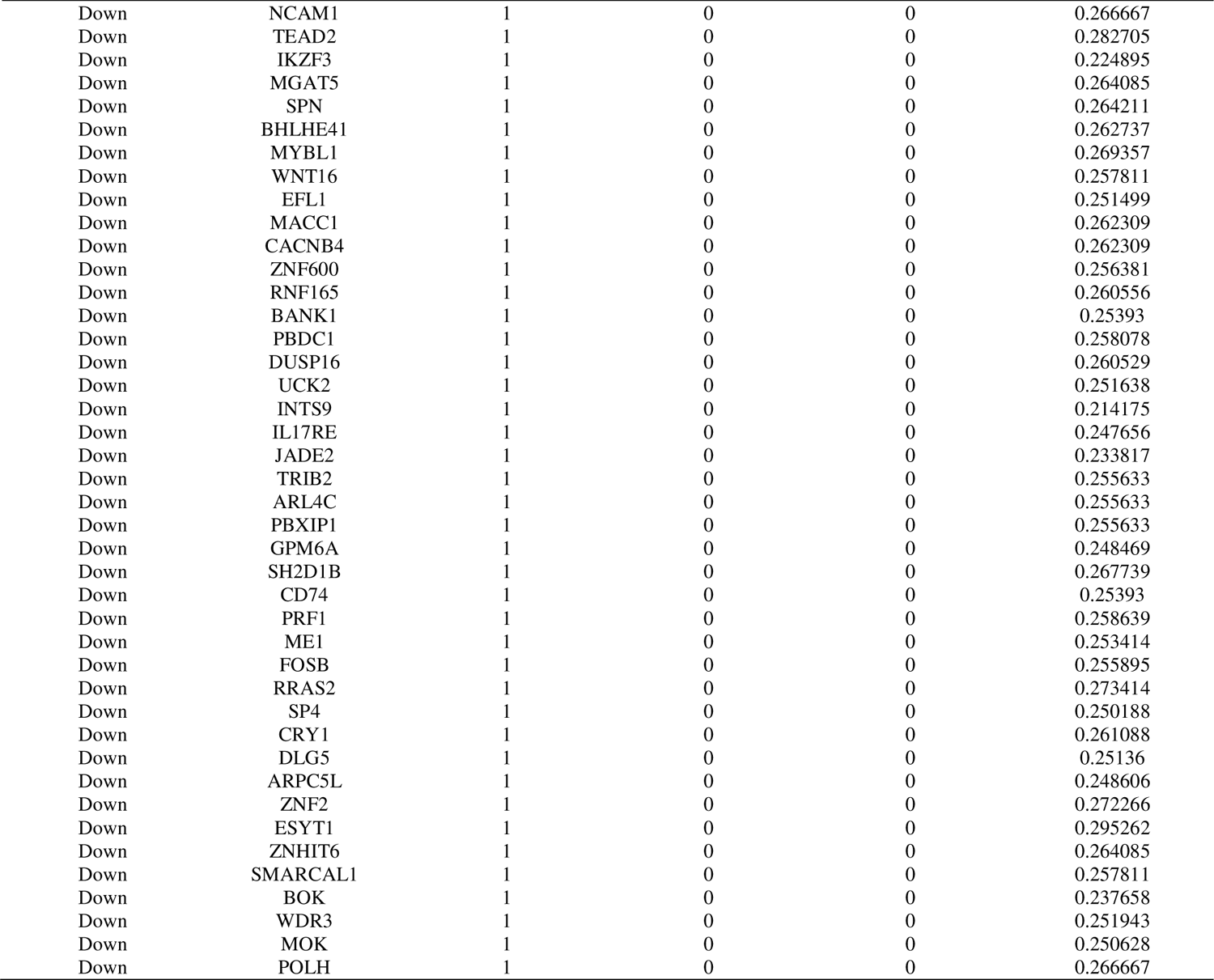
Topology table for up and down regulated genes

### miRNA-hub gene regulatory network construction

The miRNA-hub gene regulatory network was constructed and shown in Fig.5. In this network, there are nodes 2593 (miRNAs: 2278; hub genes: 315) and 17753 edges. In detail, MAPK14 regulates 88 miRNAs (ex; hsa-mir-6757-3p); LMNB1 regulates 77 miRNAs (ex; hsa-mir-4753-5p); CEBPB regulates 54 miRNAs (ex; hsa-mir-382-3p); UBE2D1 regulates 51 miRNAs (ex; hsa-mir-571); SRPK1 regulates 35 miRNAs (ex; hsa-mir-6796-3p); TUBB regulates 96 miRNAs (ex; hsa-mir-208b-3p); HSPA8 regulates 86 miRNAs (ex; hsa-mir-4436b-5p); XRCC6 regulates 84 miRNAs (ex; hsa-mir-506-5p); PARP1 regulates 78 miRNAs (ex; hsa-mir-519b-3p); HSP90AB1 regulates 74 miRNAs (ex; hsa-mir-3144-5p) and are listed in Table 5.

**Fig. 5.**
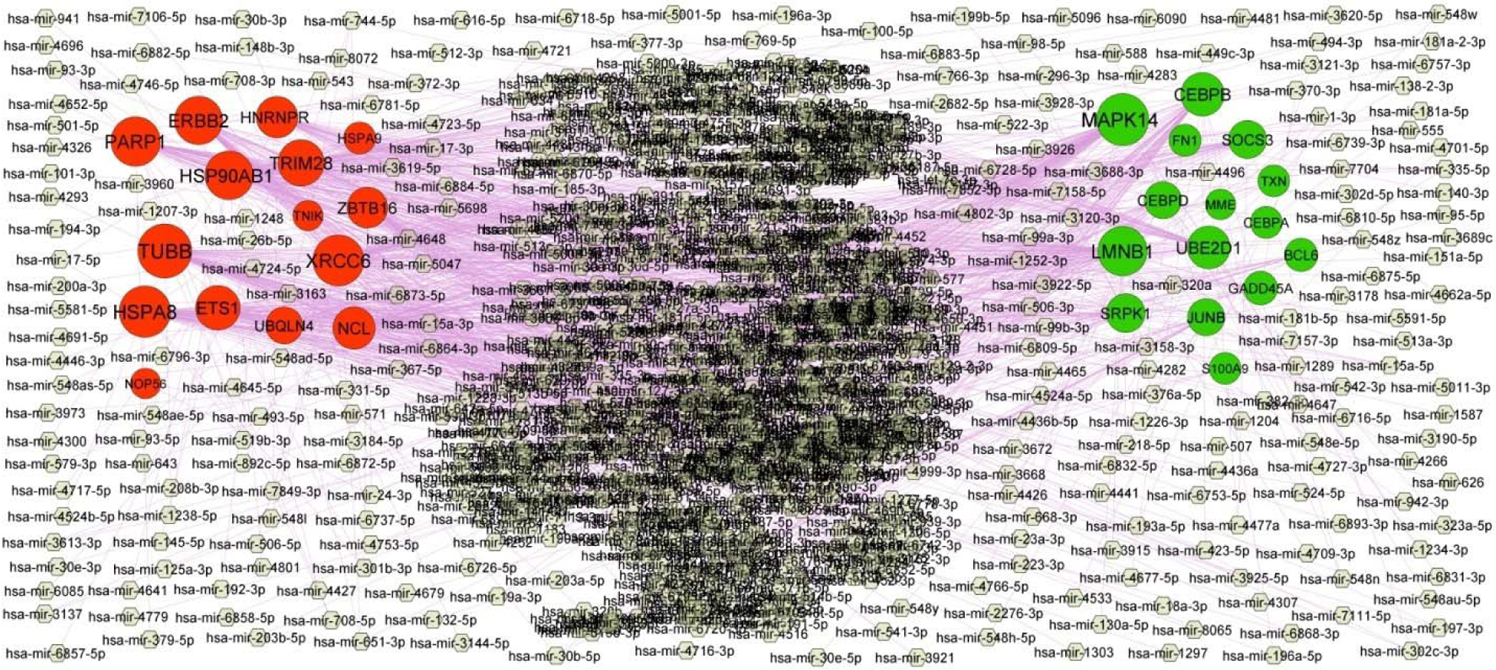
Target gene - miRNA regulatory network between target genes. The gray color diamond nodes represent the key miRNAs; up regulated genes are marked in green; down regulated genes are marked in red.

**Table 5.** miRNA - target gene and TF - target gene interaction

| Regulation | Target Genes | Degree | MicroRNA | Regulation | Target Genes | Degree | TF |
| --- | --- | --- | --- | --- | --- | --- | --- |
| Up | MAPK14 | 88 | hsa-mir-6757-3p | Up | BCL6 | 17 | USF2 |
| Up | LMNB1 | 77 | hsa-mir-4753-5p | Up | CEBPB | 15 | TFAP2C |
| Up | CEBPB | 54 | hsa-mir-382-3p | Up | JUNB | 15 | RELA |
| Up | UBE2D1 | 51 | hsa-mir-571 | Up | FN1 | 14 | STAT1 |
| Up | SRPK1 | 35 | hsa-mir-6796-3p | Up | LMNB1 | 12 | FOXF2 |
| Up | SOCS3 | 32 | hsa-mir-19a-3p | Up | GADD45A | 11 | YY1 |
| Up | CEBPD | 28 | hsa-mir-98-5p | Up | MAPK14 | 10 | FOS |
| Up | JUNB | 28 | hsa-mir-3178 | Up | CEBPA | 10 | NFKB1 |
| Up | GADD45A | 15 | hsa-mir-3613-3p | Up | MME | 9 | ARID3A |
| Up | BCL6 | 13 | hsa-mir-30b-5p | Up | SOCS3 | 9 | TEAD1 |
| Up | FN1 | 8 | hsa-mir-140-3p | Up | UBE2D1 | 9 | POU2F2 |
| Up | CEBPA | 8 | hsa-mir-193a-5p | Up | CEBPD | 7 | CREB1 |
| Up | S100A9 | 7 | hsa-mir-4679 | Up | TXN | 4 | RUNX1 |
| Up | TXN | 7 | hsa-mir-6882-5p | Up | SRPK1 | 3 | IRF2 |
| Up | MME | 2 | hsa-mir-26b-5p | Up | S100A9 | 2 | FOXC1 |
| Down | TUBB | 96 | hsa-mir-208b-3p | Down | ERBB2 | 17 | JUN |
| Down | HSPA8 | 86 | hsa-mir-4436b-5p | Down | ETS1 | 16 | ELK4 |
| Down | XRCC6 | 84 | hsa-mir-506-5p | Down | HSPA8 | 12 | SRF |
| Down | PARP1 | 78 | hsa-mir-519b-3p | Down | ZBTB16 | 9 | KLF5 |
| Down | HSP90AB1 | 74 | hsa-mir-3144-5p | Down | TRIM28 | 9 | TP53 |
| Down | ERBB2 | 73 | hsa-mir-4441 | Down | UBQLN4 | 9 | E2F1 |
| Down | TRIM28 | 65 | hsa-mir-6737-5p | Down | XRCC6 | 8 | NRF1 |
| Down | ETS1 | 58 | hsa-mir-543 | Down | TUBB | 7 | TEAD1 |
| Down | NCL | 49 | hsa-mir-4802-3p | Down | HSPA9 | 7 | GATA2 |
| Down | HNRNPR | 46 | hsa-mir-93-5p | Down | TNIK | 7 | PRDM1 |
| Down | ZBTB16 | 43 | hsa-mir-301b-3p | Down | HSP90AB1 | 6 | ZNF354C |
| Down | UBQLN4 | 28 | hsa-mir-1207-3p | Down | NCL | 5 | GATA3 |
| Down | HSPA9 | 8 | hsa-mir-320a | Down | PARP1 | 4 | BRCA1 |
| Down | NOP56 | 2 | hsa-mir-24-3p | Down | NOP56 | 4 | SREBF1 |
| Down | TNIK | 1 | hsa-mir-335-5p | Down | HNRNPR | 4 | MAX |

### TF-hub gene regulatory network construction

The TF-hub gene regulatory network was constructed and shown in Fig.6. In this network, there are nodes 393 (TFs: 85; hub genes: 308) and 2544 edges. In detail, BCL6 regulates 17 TFs (ex; USF2); CEBPB regulates 15 TFs (ex; TFAP2C); JUNB regulates 15 TFs (ex; RELA); FN1 regulates 14 TFs (ex; STAT1); LMNB1 regulates 12 TFs (ex; FOXF2); ERBB2 regulates 17 TFs (ex; JUN); ETS1 regulates 16 TFs (ex; ELK4); HSPA8 regulates 12 TFs (ex; SRF); ZBTB16 regulates 9 TFs (ex; KLF5); TRIM28 regulates 9 TFs (ex; TP53) and are listed in Table 5.

**Fig. 6.**
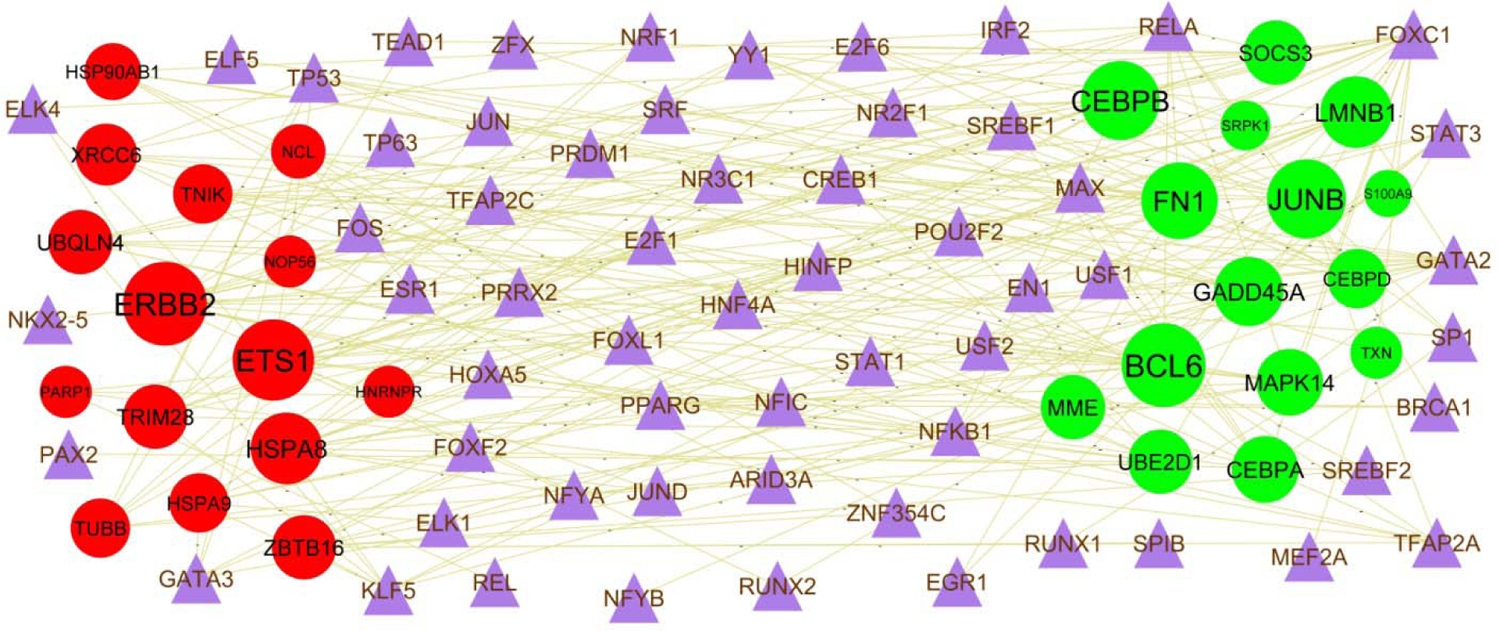
Target gene - TF regulatory network between target genes. The violet color triangle nodes represent the key TFs; up regulated genes are marked in green; down regulated genes are marked in red.

### Receiver operating characteristic curve (ROC) analysis

ROC curve analysis using “pROC” packages was performed to calculate the capacity of hub genes to distinguish CF from normal control. FN1, UBE2D1, SRPK1, MAPK14, CEBPB, HSP90AB1, HSPA8, XRCC6, NCL and PARP1 all exhibited excellent diagnostic efficiency (AUC > 0.8) (Fig.7).

**Fig. 7.**
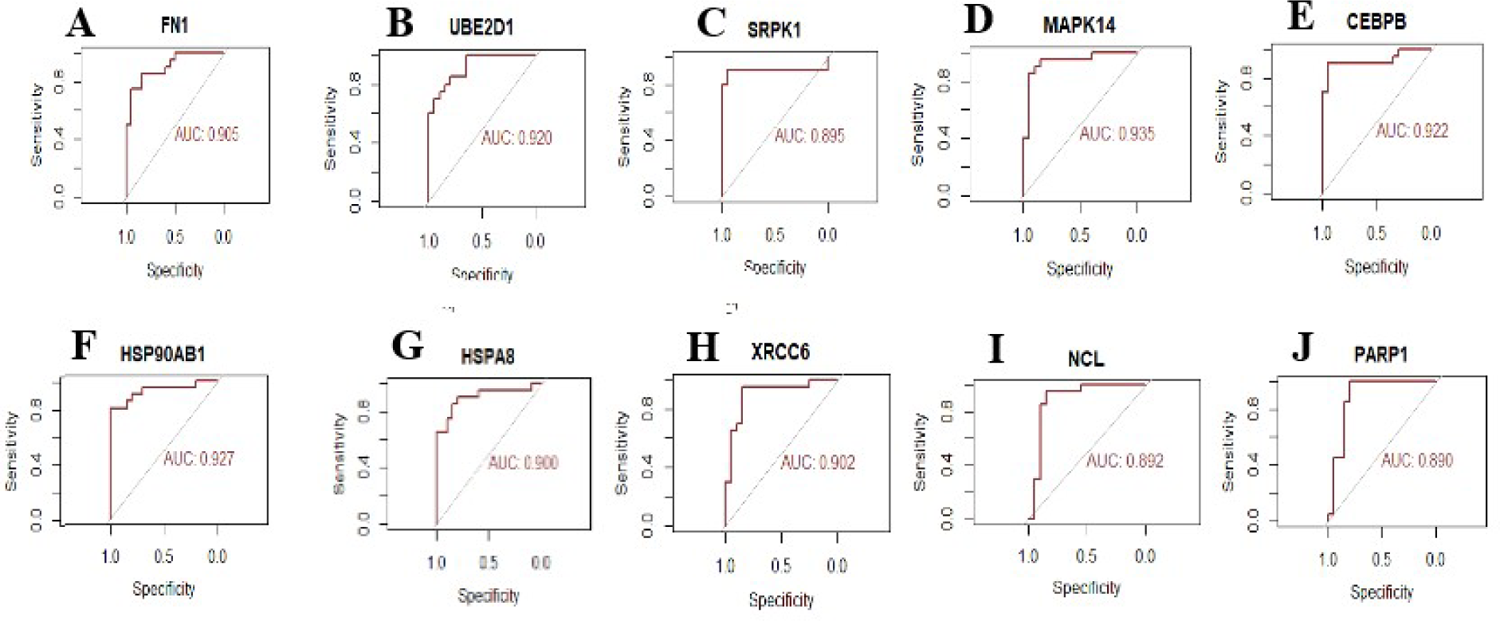
ROC curve analyses of hub genes. A) FN1 B) UBE2D1 C) SRPK1 D) MAPK14 E) CEBPB F) HSP90AB1 G) HSPA8 H) XRCC6 I) NCL J) PARP1

## Discussion

Although various relevant investigation of CF has been performed, early diagnosis, efficacy of treatment and prognosis for CF remain poorly resolved. For diagnosis and treatment, it is necessary to further understand the molecular mechanisms resulting in progression and advancement. Due to the advancement of NGS technology, the genetic modifications due to disease advancement can be detected, indicating gene targets for diagnosis, therapy and prognosis of specific diseases. In this present investigation, we downloaded the GSE136371 NGS dataset from GEO, and a total of 917 DEGs (479 up regulated genes and 438 down regulated genes) between CF and normal control samples were identified by using bioinformatics tools. CHGA (chromogranin A) [48] and IL23R [49] have been reported to be positively correlated with gastrointestinal complications. Recent studies have shown that the CHGA (chromogranin A) [50], DLK1 [51], CHGB (chromogranin B) [52], NR2E1 [53], DBH (dopamine beta-hydroxylase) [54], CYP17A1 [55], IL23R [56] and GLRA3 [57] mediates diabetes mellitus. CHGA (chromogranin A) [58], CD177 [59], IL23R [56] and LGALS9B [60] have been proposed as biomarkers for arthritis. Enterina et al [61] found that DLK1 positively correlate with pulmonary complications. Altered expression of DLK1 [62], DBH (dopamine beta-hydroxylase) [63] and CYP17A1 [64] have previously been shown to promote depression and anxiety. DLK1 [65] and IL23R [66] were shown to participate in facilitating the osteoporosis. DBH (dopamine beta-hydroxylase) [67] has been reported to enhance CF development. Previous reports have demonstrated that DBH (dopamine beta-hydroxylase) [68], CYP17A1 [69], CD177 [70], IL23R [71] and LGALS9B [72] are associated with liver diseases. Studies had shown that IL23R [73] was altered expressed in patients with endobronchial airway infection. Highly significant DEGs are the might be main causes of CF and CF associated complications.

The GO and REACTOME pathway enrichment analysis revealed that the DEGs were mainly in the progression of CF. Previous studies had shown that pathways include neutrophil degranulation [74], immune system [75], extracellular matrix organization [76], toll-like receptor cascades [77], cytokine signaling in immune system [78], metabolism of RNA [79] and gene expression [80] were involved in the progression of CF. NTRK3 [81], MMP9 [82], IGF2 [83], SOCS3 [84], IGFBP2 [85], SLC1A3 [86], NOS1AP [87], HP (haptoglobin) [88], TLR5 [89], ARG1 [90], NRG1 [91], RGS5 [92], LCN2 [93], TSPO (translocator protein) [94], PLAU (plasminogen activator, urokinase) [95], MME (membrane metalloendopeptidase) [96], BST1 [97], NR6A1 [98], TLR4 [99], NRN1 [100], ABCA13 [101], CTSD (cathepsin D) [102], NCAM1 [103], SIGMAR1 [104], CNR2 [105], CCR4 [106], ABCB1 [107], TIMELESS (timeless circadian regulator) [108], IRF8 [109], TXNIP (thioredoxin interacting protein) [110], SUV39H1 [111], CRY1 [112], CRY2 [113], PDCD4 [114] and MGAT5 [115] played an important role in depression and anxiety. Accumulating evidence has demonstrated that MMP9 [116], S100A12 [117], HP (haptoglobin) [118], TLR5 [119], NRG1 [120], PLAU (plasminogen activator, urokinase) [121], SLC11A1 [122], AQP9 [123], CHIT1 [124], TLR4 [125], SLC26A8 [126], CTSD (cathepsin D) [127], SERPINB1 [128], FASLG (Fas ligand) [129], SLC4A4 [130], AQP3 [131] and IRF8 [132] appears to be constitutively associated with **CF**. MMP9 [133], S100A12 [134], SOCS3 [135], MMP8 [136], CRISP3 [137], S100A9 [138], RETN (resistin) [139], IL1RN [140], TLR4 [141], GGT1 [142], CTSD (cathepsin D) [143], FASLG (Fas ligand) [144], CCR4 [145], FCRL3 [146] and ABCF1 [147] were revealed and regarded as diagnostic biomarker in pancreatitis. Accumulating evidence shows that MMP9 [133], S100A12 [148], HP (haptoglobin) [149], OSM (oncostatin M) [150], PRDM5 [151], TSPO (translocator protein) [152], IL18RAP [153], ADAMDEC1 [154], IL1RN [155], SCN9A [156], FADS2 [157], NCF4 [158], SERPINB1 [159], TNFRSF13B [160], IL2RB [161], DLG5 [162], FASLG (Fas ligand) [163], GPR68 [164], IL2RA [161], TCF4 [165], TAGAP (T cell activation RhoGTPase activating protein) [166], ABCB1 [167], FCRL3 [168], ITGB7 [169], PTGER4 [170] and TRAF3IP2 [171] are altered expression in gastrointestinal complications. Recent studies reported that MMP9 [133], S100A12 [172], IGF2 [173], GPR84 [174], SOCS3 [175], IGFBP2 [176], HP (haptoglobin) [177], MMP8 [178], OSM (oncostatin M) [179], TLR5 [180], S100A9 [181], PLIN5 [182], NRG1 [183], LCN2 [184], TSPO (translocator protein) [185], PLAU (plasminogen activator, urokinase) [186], PPBP (pro-platelet basic protein) [187], UPP1 [188], ALOX5 [189], IL1RN [190], CYP1B1 [191], DGAT2 [192], MSRA (methionine sulfoxidereductase A) [193], ADAM9 [194], CPEB4 [195], IRS2 [196], FADS2 [197], SLC22A4 [198], PFKFB3 [199], CTSD (cathepsin D) [200], ADAM12 [201], CD80 [202], ERBB2 [203], FASLG (Fas ligand) [204], CNR2 [205], SOCS2 [206], ABCB1 [207], CD74 [208], FCRL3 [209], PARP1 [210], TXNIP (thioredoxin interacting protein) [211], TULP3 [212], HSPA8 [213], TNIK (TRAF2 and NCK interacting kinase) [214], PDCD4 [215] and USP11 [216] are altered expression in the liver diseases. MMP9 [217], S100A12 [218], CBS (cystathionine beta-synthase) [219], SOCS3 [220], IGFBP2 [221], MMP8 [222], OSM (oncostatin M) [223], TLR5 [224], S100A8 [225], PDE6H [226], CEBPB (CCAAT enhancer binding protein beta) [227], PLAU (plasminogen activator, urokinase) [228], BCL6 [229], CD55 [230], ADAMDEC1 [231], LTBP2 [232], TGFA (transforming growth factor alpha) [233], IL1RN [234], ALDH1A2 [235], PLXNC1 [236], MSRA (methionine sulfoxidereductase A) [237], TLR4 [238], SIGLEC9 [239], CPEB4 [240], IL17RA [241], IRS2 [242], ETS2 [243], PFKFB3 [244], NQO2 [245], CTSD (cathepsin D) [246], SERPINB1 [247], ADAMTS1 [248], SOX5 [249], FASLG (Fas ligand) [250], PTPN13 [251], CCR4 [252], SOCS2 [253], PTCH1 [254], ABCB1 [255], CD81 [256], PTGER4 [257], TXNIP (thioredoxin interacting protein) [258], DUSP16 [259], DKC1 [260] and ST6GAL1 [261] are associated with the risk of pulmonary complications. Previous studies have described the presence of MMP9 [262], S100A12 [263], NOS1AP [264], MMP8 [265], OSM (oncostatin M) [266], CAMP (cathelicidin antimicrobial peptide) [267], LCN2 [268], ADAMDEC1 [269], IL1RN [270], TLR4 [271] and TXNIP (thioredoxin interacting protein) [272] as biomarkers of sinusitis. MMP9 [262], IGF2 [273], SOCS3 [274], MMP8 [265], OSM (oncostatin M) [266], PLAU (plasminogen activator, urokinase) [275], ADAMDEC1 [269], IL1RN [270], TLR4 [271], CD80 [276] and TXNIP (thioredoxin interacting protein) [272] make great contributions to the progression of nasal polyps. MMP9 [277], S100A12 [278], IGF2 [279], GPR84 [280], SOCS3 [281], ANXA3 [282], IGFBP2 [283], NOS1AP [284], G0S2 [285], HP (haptoglobin) [286], MMP8 [287], SLC6A19 [288], OSM (oncostatin M) [289], S100A8 [290], FFAR3 [291], ARG1 [292], ADM (adrenomedullin) [293], S100A9 [294], NRG1 [295], IL1R1 [296], IL4R [297], TSPO (translocator protein) [298], TXN (thioredoxin) [299], PLAU (plasminogen activator, urokinase) [300], SLC11A1 [301], TRPM2 [302], RETN (resistin) [303], NLRC4 [304], CD55 [305], ALOX5AP [306], GPR160 [307], TRPM6 [308], ALOX5 [309], IL1RN [310], ADM2 [311], DGAT2 [312], TLR4 [313], ENTPD1 [314], GRB10 [315], IL17RA [316], IRS2 [317], MGAM (maltase-glucoamylase) [318], FADS2 [319], SLC22A4 [320], KCNJ15 [321], PPP1R3B [322], CTSD (cathepsin D) [323], SERPINB11 [324], COL4A3 [325], ADAM12 [326], SOX5 [327], CD80 [328], ERBB2 [329], POU2AF1 [330], FASLG (Fas ligand) [331], SOX13 [332], BANK1 [333], IL2RA [334], AQP3 [335], CCR4 [336], CD74 [337], FCRL3 [338], ITGB7 [339], PARP1 [340], TXNIP (thioredoxin interacting protein) [341], SUV39H1 [342], HSPA8 [343], ETS1 [344], ELP5 [345] and PDCD4 [346] might be important for diabetes mellitus progression. Previous research found that MMP9 [347], CBS (cystathionine beta-synthase) [348], IGF2 [349], IGFBP2 [350], HP (haptoglobin) [351], MAPK14 [352], LRG1 [353], LCN2 [354], SBNO2 [355], ITGA1 [356], IL1R2 [357], IL1RN [358], TLR4 [359], IRS2 [360], CORIN (corin, serine peptidase) [361], WNT16 [362], CD80 [363], FASLG (Fas ligand) [364], CXXC5 [365], CNR2 [366], TCF4 [367], PTCH1 [368], TXNIP (thioredoxin interacting protein) [369], TRAF3IP2 [370], ETS1 [371] and ARHGEF3 [372] are strongly associated with osteoporosis. MMP9 [373], S100A12 [374], IGF2 [375], GPR84 [376], SOCS3 [377], CEBPA (CCAAT enhancer binding protein alpha) [378], PGLYRP1 [379], HP (haptoglobin) [380], MMP8 [381], OSM (oncostatin M) [382], TLR5 [383], S100A8 [384], ARG1 [385], S100A9 [386], BCL2A1 [387], IRAK3 [388], ST3GAL4 [389], IL1R1 [390], MAPK14 [391], IL4R [392], PADI4 [393], LCN2 [394], CEBPB (CCAAT enhancer binding protein beta) [395], TSPO (translocator protein) [396], IL18RAP [397], PLAU (plasminogen activator, urokinase) [398], CDA (cytidinedeaminase) [399], SLC11A1 [400], RETN (resistin) [401], GADD45A [402], BCL6 [403], CD55 [404], THBS1 [405], IL1R2 [406], TGFA (transforming growth factor alpha) [407], IL1RN [408], ALDH1A2 [409], SLC8A3 [410], HMGB2 [411], NFIL3 [412], MSRA (methionine sulfoxidereductase A) [413], ITGAM (integrin subunit alpha M) [414], TLR4 [415], SIGLEC9 [416], GRB10 [417], IL17RA [241], SLC2A3 [418], GCA (grancalcin) [419], SLC22A4 [420], PLB1 [421], SEMA4A [422], STEAP4 [423], NCF4 [424], ETS2 [425], ENHO (energy homeostasis associated) [426], ADAM12 [427], SOX5 [428], IL2RB [429], WNT16 [430], DLG5 [431], CD80 [432], KLRB1 [433], TBX21 [434], CD1C [435], ERBB2 [436], HLA-DOA [437], CD79A [438], POU2AF1 [439], FASLG (Fas ligand) [440], NMUR1 [441], BANK1 [442], IL2RA [443], CNR2 [444], CCR4 [445], TCF4 [446], CARD11 [447], TAGAP (T cell activation RhoGTPase activating protein) [448], PTCH1 [449], ABCB1 [450], TEF (TEF transcription factor, PAR bZIP family member) [451], CD247 [452], CD81 [453], CD74 [454], FCRL3 [168], PTGER4 [455], PARP1 [456], SHMT2 [457], TXNIP (thioredoxin interacting protein) [458], CABIN1 [459], SLAMF6 [460], TRAF3IP2 [461], ETS1 [462], USP5 [463] and ST6GAL1 [464] are a known biomarkers for the onset of arthritis. Previous research found that S100A12 [465], SOCS3 [466], ITGB4 [467], HP (haptoglobin) [468], OSM (oncostatin M) [469], TLR5 [470], LCN2 [471], SLC11A1 [472], NLRC4 [473], CD55 [474], TLR4 [475], CKAP4 [476], PFKFB3 [477], SERPINB1 [478] and NMUR1 [479] are strongly associated with endobronchial airway infection. Therefore, these GO terms and signaling pathways are most likely to be essential in the advancement of CF and CF associated complications. More investigation is required to identify all the DEGs in CF and CF associated complications.

In this investigation, FN1, UBE2D1, SRPK1, MAPK14, CEBPB, FCAR (Fc fragment of IgA receptor), S100A9, CEBPA, CEBPD (CCAAT enhancer binding protein delta), ALOX5AP, HSP90AB1, HSPA8, XRCC6, NCL (nucleolin), PARP1, NOP56, NOP2 and NOLC1 are considered to be the hub genes in the PPI network and modules. FN1, UBE2D1, SRPK1, FCAR, CEBPD, HSP90AB1, XRCC6, NCL, NOP56, NOP2 and NOLC1 are new biomarkers for the occurrence of CF and CF associated complications. Other hub genes were might be involved in the pathogenesis of CF and CF associated complications and believe that the specific mechanism is worthy of further investigation.

Finally, by constructing miRNA-hub gene regulatory network and TF-hub gene regulatory network, we found that hub genes, miRNAs and TFs might involved in CF and CF associated complications. USF2 [480], RELA [481], STAT1 [482], KLF5 [483] and TP53 [484] were closely related to arthritis. RELA [485] is believed to be related to the occurrence of pancreatitis. Studies have shown that STAT1 [486] is essential for the development of CF. Recent studies have proposed that the STAT1 [487] and TP53 [488] are associated with depression and anxiety. A study revealed that STAT1 [489], and KLF5 [490] were significantly regulated in liver diseases. STAT1 [491] and SRF (Serum response factor) [492] expression has a role in pulmonary complications. STAT1 [493] level was significantly altered in nasal polyps. Studies had shown that STAT1 [494], KLF5 [495] and TP53 [496] expression was associated with diabetes mellitus. STAT1 [497] and TP53 [498] were a diagnostic markers of osteoporosis. and could be used as therapeutic targets.There is no research showing that LMNB1, JUNB, TUBB, ZBTB16, TRIM28, hsa-mir-6757-3p, hsa-mir-4753-5p, hsa-mir-382-3p, hsa-mir-571, hsa-mir-6796-3p, hsa-mir-208b-3p, hsa-mir-4436b-5p, hsa-mir-506-5p, hsa-mir-519b-3p, hsa-mir-3144-5p, TFAP2C, FOXF2, JUN and ELK4 are related to CF and CF associated complications, and it is a new biomarkers for the occurrence of CF and CF associated complications. These findings, together, suggest that these biomarkers might lead to CF and CF associated complications, although further research is required.

In conclusion, the present investigation identified key genes and pathways which might be involved in CF and CF associated complications progression through the integrated analysis of NGS dataset. These results might contribute to a better understanding of the molecular mechanisms which underlie CF and CF associated complications and provides a series of potential biomarkers. However, further investigations are required to verify the findings of the current investigation. These findings significantly improve the understanding of the cause and underlying molecular events in CF and CF associated complications, and the candidate genes and pathways could be used as therapeutic targets.

## Acknowledgement

I thank Albertina DE SARIO, Montpellier University, IURC, Génétique de Maladies Rares, Montpellier, France, very much, the author who deposited their NGS dataset GSE136371, into the public GEO database.

## Ethical approval

This article does not contain any studies with human participants or animals performed by any of the authors.

## Informed consent

No informed consent because this study does not contain human or animals participants.

## Availability of data and materials

The datasets supporting the conclusions of this article are available in the GEO (Gene Expression Omnibus) (https://www.ncbi.nlm.nih.gov/geo/) repository. [(GSE136371) https://www.ncbi.nlm.nih.gov/geo/query/acc.cgi?acc=GSE136371]

## Consent for publication

Not applicable.

## Author Contributions

B. V. - Writing original draft, and review and editing

C. V. - Software and investigation

## Authors

Basavaraj Vastrad ORCID ID: 0000-0003-2202-7637

Chanabasayya Vastrad ORCID ID: 0000-0003-3615-4450

